# Integration of differential gene expression with weighted gene correlation network analysis identifies genes whose expression is remodeled throughout physiological aging in mouse tissues

**DOI:** 10.1101/2021.02.18.431793

**Authors:** Margarida Ferreira, Stephany Francisco, Ana R. Soares, Ana Nobre, Miguel Pinheiro, Andreia Reis, Sonya Neto, Ana João Rodrigues, Nuno Sousa, Gabriela Moura, Manuel A. S. Santos

**Affiliations:** Institute of Biomedicine – iBiMED, Department of Medical Sciences, University of Aveiro, 3810-193 Aveiro, Portugal; Life and Health Sciences Research Institute (ICVS), School of Medicine, University of Minho, 4710-057 Braga, Portugal; ICVS/3B’s–PT Government Associate Laboratory, Braga/Guimarães, Portugal

**Keywords:** Aging, transcriptome, *Mus musculus*, DESeq2, WGCNA

## Abstract

Gene expression alterations occur in all mouse tissues during aging, but recent works highlight minor rather than major dysregulation amplitude for most genes, questioning whether differentially expressed genes on their own provide deep insight into aging biology. To clarify this issue, we have combined differential gene expression with weighted gene correlation network analysis (WGCNA) to identify expression signatures accounting for the pairwise relations between gene expression profiles and the cumulative effect of genes with small fold-changes during aging in the brain, heart, liver, skeletal muscle, and pancreas of C57BL/6 mice. Functional enrichment analysis of the overlap of genes identified in both approaches showed that immunity-related responses, mitochondrial energy metabolism, tissue regeneration and detoxification are prominently altered in the brain, heart, muscle, and liver, respectively, reflecting an age-related global loss of tissue function. While data showed little overlap among the age-dysregulated genes between tissues, aging triggered common biological processes in distinct tissues, particularly proteostasis-related pathways, which we highlight as important features of murine tissue physiological aging.

## Introduction

Gene expression alterations occurring throughout the lifespan have been described for a multitude of species, organs, and cell types [1–10]. The most commonly reported age-related dysregulations involve the immune system [9,11–13] where inflammatory response genes are upregulated even in the absence of pathogen infection [5,6,9,11,14–19]. Energy metabolism, redox homeostasis, and mitochondrial function alterations are also frequently observed in age-related studies [6,9,11,15–18,20], particularly the downregulation of genes encoding mitochondrial ribosomal proteins and components of the electron transport chain [5,11,14–16,18], protein synthesis machinery [5,11,17], developmental and cell differentiation genes [9,11,19], and extracellular matrix components [6,14–16]. Up-regulated genes are associated with the stress response and DNA repair [5,6,9,11,14,16–18], RNA processing [11,12,17] and cell cycle arrest [5,16,21]. Despite this, the existence of specific genetic signatures of aging continue to be a matter of debate as gene regulation is mostly tissue-specific [5–7,15,20,22–25], but also because there is focus on comparisons between young and old individuals without much consideration of the dynamics of gene expression throughout the lifespan. Nonetheless, there is some evidence in humans and animal models shedding light on these dynamics. As an example, a marked shift in mRNA and microRNA expression (namely in regulators of genes involved in nervous system development and function, neurological diseases, and cell-cell signaling) has been reported to occur at around age 20 in the human prefrontal cortex [26, 27]. Less striking alterations are reported in the same brain region between 30-60 years [27, 28], entailing genes related to the synapse, fatty acid metabolism, purine nucleotide binding, ubiquitin proteolysis, channel activity, translation, DNA damage response, transcriptional activation, and neuronal function [29]. Late middle-age and early old-age shifts have also been described in human peripheral blood leukocytes for genes involved in cancer, hematological and immunological diseases, cell-mediated immune response and signaling pathways [30], and in the human brain and muscle for both coding and non-coding RNAs pertaining to longevity pathways [31].

In animal models, similar findings have been reported. In a study across 11 rat organs, the most frequent changes in gene expression occurred at around 6 and 21 months [32], proposed to be equivalent to middle-age in humans [33]. In a similar study across 17 mouse tissues, shift points of gene expression trajectories have been identified at around 6 months for extracellular matrix genes, 10 months for mitochondrial genes, 12 months for genes encoding heat shock proteins, and at around 15 months for immune response genes [6]. Nonetheless, evidence also suggests tissue-specific turning points in gene expression profiles [6,20,34]. For example, immune response gene expression was found to change in the mouse kidney between 13 and 20 months, in line with the previously described organismal trend, whereas in the spleen and lung this shift occurs later in life, at around 26 months [34].

However, the previous studies have mainly focused on differential gene expression to characterize the aging transcriptome and ignored the pairwise relationships between gene expression profiles, which may underlie commonly altered pathways and regulatory mechanisms with age. The cumulative effect of co-expressed genes with small fold-changes has also been disregarded, as most of these studies usually select genes with at least two-fold expression changes. Therefore, in this work, we combined differential gene expression profiling with weighted gene correlation network analysis of publicly available mouse RNA sequencing (RNA-Seq) data (GSE132040) [6,35,36] in order to: 1) identify clusters of genes significantly correlated with aging in different mouse tissues, 2) establish their trajectories from mature adulthood to old age, 3) identify the time point of the shift in gene expression profile, and 4) evaluate the biological processes (BPs) associated with the gene dysregulations. The data showed tissue-specific age-related alterations and highlighted gene expression in the pancreas as being largely unaffected across the lifespan. We also found that genes involved in lipid metabolism start to be differentially expressed relatively early and continue to exhibit altered expression until old age. Different onsets of gene dysregulation were identified, demonstrating an asynchronous impairment of gene expression with age. Gene Ontology (GO) biological processes’ over-representation analysis revealed immunity-related responses, mitochondrial energy metabolism, regeneration, and detoxification as the most prominently altered processes in the brain, heart, muscle and liver, respectively, which may reflect the global loss of organ function, confirming previous reports. Despite little overlap in age-dysregulated genes between tissues, a comparison at the level of dysregulated processes revealed inter-tissue commonalities, such as alterations in proteostasis-associated processes. We propose that the genes involved in these processes are important players of murine physiological aging, especially in the muscle, brain, and liver.

## Results

### Modest tissue-specific changes in gene expression are observed across the lifespan

In this study we used a publicly available transcriptomic dataset of aging mice, from which we selected a subset of samples from the brain, heart, muscle, liver, and pancreas, from 3 to 27 month-old male and female mice (*see Methods – Dataset characterization*).

In order to obtain a global characterization of the mouse aging transcriptome and, in particular, to understand which of the main known sources of genetic variation (tissue, age, and sex) is responsible for the highest percentage of sample segregation, we performed a principal component analysis (PCA) of all samples based on variance stabilizing transformation (VST)-normalized read counts. Principal components were calculated based on the 500 most variable genes as they are expected to capture the greatest variability between samples (*see Methods - Differential gene expression analysis*). We observed that, based on VST-normalized gene expression values, samples tend to cluster by tissue, which is in line with previous observations [5–7,15,20,22–24,37], and the reason why all subsequent analyses were performed on each tissue separately (Figure 1A). Additionally, we focused on samples from the brain, heart, muscle, liver, and pancreas, as these tissues are associated with and well-characterized in age-related diseases [38–42]. When considering gene expression variation in each tissue independently, we observed that sex, rather than age, is responsible for most of the between-sample variability and adjusted for its effect on gene expression by adding it as a co-variable in the model (Supplemental Figure S1; see *Methods – Differential gene expression analysis*).

**Figure 1.**
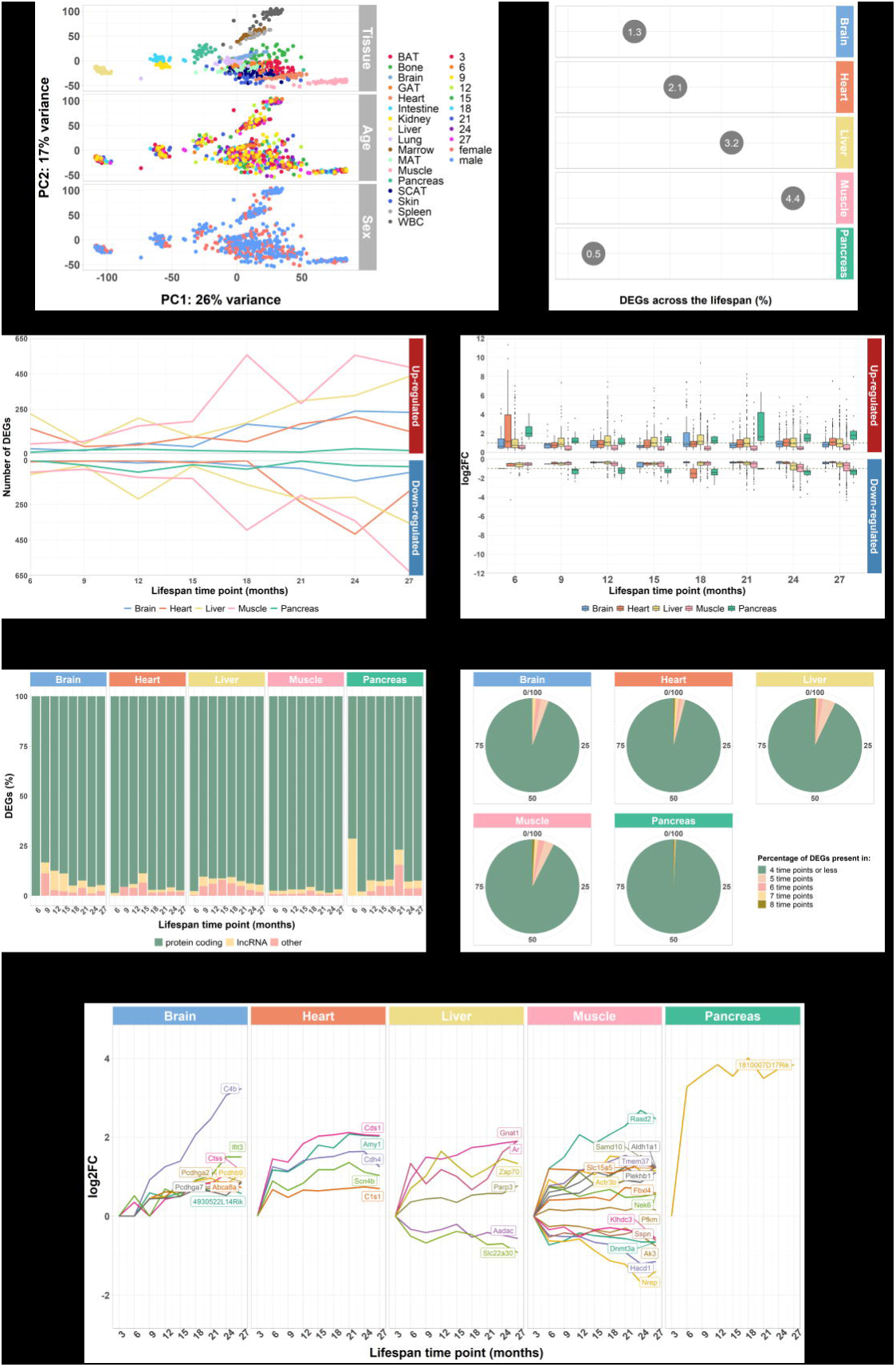
Whole-transcriptome characterization of different mouse tissues throughout the lifespan by pairwise differential expression analysis against a 3-month baseline. **A)** PCA of all tissues performed on VST-normalized read counts of the 500 most variable genes and colored by all known effects highlights type of tissue as the main contributor to sample segregation. **B)** Percentage of DEGs across the lifespan for the mouse brain, heart, liver, muscle, and pancreas, highlighting the last two tissues as the ones with higher and lower dysregulation, respectively. The displayed values correspond to the percentages of the total number of DEGs found in each tissue relative to the initial number of genes (50735). Total number of DEGs is the sum of the number of DEGs per pairwise comparison (differential expressed genes per time point against the 3-month baseline); in the case of duplicate gene IDs, only one was considered). **C)** Line plot depicting the amplitude of gene expression alterations across the lifespan for the selected tissues reveals a progressive increase in DEGs with increasing age. Line breaks correspond to the number of DEGs in each time point against the baseline (3 months). Positive values relate to up-regulated genes whereas negative values match up to down-regulated genes. **D)** Boxplots showing the distribution of log2FC values per tissue and time point (against 3-month baseline). The grey, dashed line represents log2FC = 1 and log2FC = -1 (fold-changes of 2 and 0.5, respectively). log2FC > 1 indicates more than doubling the expression, whereas log2FC < -1 points to half or less of the expression. **E)** DEG biotype distribution per tissue and time point (against the 3-month baseline). Biotype nomenclature based on Ensembl annotation. **F)** Percentage of genes differentially expressed in over half of the evaluated lifespan (5, 6, 7, and 8 time points) or in half (4 time points) or less (3, 2, and 1 time points). **G)** Trajectories across the lifespan of the TDEGs per tissue (brain: 7 time points; heart, liver, muscle, and pancreas: 8 time points).

As a first approach to establish age-regulated genes, we performed differential expression between each lifespan time point (6, 9, 12, 15, 18, 21, 24 and 27 months) relative to the reference time point (3 months) on a total of 50735 genes (post-filtering, see *Methods – Differential gene expression analysis*) and selected the differentially expressed genes (DEGs) in each pairwise comparison, and for each tissue. Next, to obtain a global view of the amplitude of gene expression dysregulation of each tissue with aging, we calculated the percentage of DEGs across the lifespan, considering each tissue’s total number of DEGs as the sum of the DEGs in each time point relative to the 3-month age reference, and keeping only unique gene IDs. We identified a total of 684, 1051, 1623, 2233, and 277 differentially expressed genes in the brain (1.3%), heart (2.1%), liver (3.2%), muscle (4.4%), and pancreas (0.5%), respectively (Figure 1B). These results highlight the pancreas and muscle with the lowest and the highest number of genes changing their expression with age, respectively. Although age-related transcriptomic alterations are globally modest in number, this is in agreement with past literature for a variety of tissues and animal models [recently reviewed in 17]. When considering the amplitude of tissue-specific DEGs per time point, we observed a global trend of increasing transcriptomic changes with increasing age, with an onset at the shift from middle- to old-age (15 to 18 months) (Figure 1C). In addition to the amplitude of gene expression dysregulation, we also evaluated the magnitude of these alterations. In general, up-regulated genes exhibit a wider range of fold-changes (FC) than the down-regulated genes (Figure 1D). Interestingly, most of the genes in the muscle exhibited very low FCs, failing to reach the commonly accepted threshold of double expression (log2 fold-change (log2FC) = 1; FC = 2), whereas in the pancreas, the majority of genes consistently more than doubled their expression in each time point when compared to the 3 months (Figure 1D, upper panel). Accordingly, most of down-regulated genes in the muscle exhibited a less than two-fold decrease in expression, while the reduction in the expression of most pancreas DEGs was more than half (Figure 1D, lower panel). Biotype assessment of the DEGs in each tissue and time point based on the Ensembl classification (see *Methods* – *Differential gene expression analysis*) showed that dysregulation mainly occurs at the protein coding level (Figure 1E). Nonetheless, changes in the expression of long non-coding RNAs may be of relevance, especially in the brain, liver, and pancreas (Figure 1E).

We then asked whether the DEGs found in each time point were specific to that comparison or if they were present in other time points, i.e., if they were pervasive throughout aging. The data showed that the vast majority of the DEGs were dysregulated in half or less of the lifespan only (Figure 1F, “4 time points or less”). We also identified the genes that were differentially expressed in the highest number of lifespan time points in each tissue and considered them to be “Top DEGs” (TDEGs) as they were dysregulated throughout most of the lifespan (Figure 1E; Table 1; Supplemental File S1). The muscle and the pancreas stood out from the rest of the tissues by exhibiting the highest (16) and the smallest (1) number of pervasively dysregulated genes, respectively. Moreover, each tissue exhibited a different set of the TDEGs, suggesting that the age-related gene expression dysregulation is mainly tissue-specific (Table 1).

**Table 1.** DEG pervasiveness across the lifespan. Per tissue list of TDEGs, number of time points in which TDEGs were found to be differentially expressed and associated GO Biological Processes. All annotated processes were retrieved from the AmiGO2 webtool [90] (Supplemental File S1).

| Tissue | Time points (no.) | TDEGs (no. and direction) |  | Genes involved | GO Biological processes |
| --- | --- | --- | --- | --- | --- |
| Brain | 7 | 8 | Up | *4930522L14Rik*, *Abca8a*, *C4b*, *Ctss*, *Ifit3*, *Pcdhb9*, *Pcdhga2*, *Pcdhga7* | Transcription regulation: *4930522L14Rik*<br>Lipid transport: *Abca8a*<br>Immune system processes: *C4b*, *Ctss*, *Ifit3*<br>Cell adhesion: *Pcdhb9*, *Pcdhga7*<br>NA: *Pcdhga2* |
|  |  | 0 | Down | NA | NA |
| Heart | 8 | 5 | Up | *Amy1*, *C1s1*, *Cdh4*, *Cds1*, *Scn4b* | Carbohydrate metabolism: *Amy1*<br>Complement system activation: *C1s1*<br>Cell adhesion: *Cdh4*<br>Lipid metabolism: *Cds1*<br>Cardiac action potential: *Scn4b* |
|  |  | 0 | Down | NA | NA |
| Liver | 8 | 4 | Up | *Ar*, *Gnat1*, *Parp3*, *Zap70* | Transcription regulation: *Ar*<br>Signal transduction: *Gnat1*<br>DNA repair: *Parp3*<br>Adaptive immune response: *Zap70* |
|  |  | 2 | Down | *Aadac*, *Slc22a30* | Triglyceride metabolism: *Aadac*<br>Anion transport: *Slc22a30* |
| Muscle | 8 | 10 | Up | *Actr3b*, *Aldh1a1*, *Fbxl4*, *Nek6*, *Pfkm*, *Plekhhb1*, *Rasd2*, *Samd10*, *Slc15a5*, *Tmem37* | Actin filament organization: *Actr3b*<br>Retinoid metabolism: *Aldh1a1*<br>Ubiquitin-dependent protein catabolism: *Fbxl4*<br>Cell cycle: *Nek6*<br>Glycogen catabolism: *Pfkm*<br>Cell differentiation: *Plekhhb1*<br>Signal transduction: *Rasd2*<br>Protein transport: *Slc15a5*<br>Ion transport: *Tmem37*<br>NA: *Samd10* |
| | | 6 | Down | *Ak3*, *Dnmt3a*, *Hacd1*, *Klhdc3*, *Nrep*, *Sspn* | Energy metabolism: *Ak3*<br>DNA methylation: *Dnmt3a*<br>Fatty acid metabolism: *Hacd1*<br>Meiotic cell cycle: *Klhdc3*<br>TGF-β receptor signaling pathway: *Nrep*<br>NA: *Sspn* |
| Pancreas | 8 | 1 | Up | *1810007D17Rik* | NA: *1810007D17Rik* |
|  |  | 0 | Down | NA | NA |

NA – not applicable/available

### Different subsets of co-expressed genes exhibit specific age-related trajectories

As a second approach to defining age-regulated genes, we performed weighted gene correlation network analysis (WGCNA) to explore the co-expression patterns of gene expression over time. Similar to the differential expression analysis, WGCNA was performed on each tissue independently, with each tissue’s co-expression network comprising a variable number of modules of positively correlated genes (Supplemental File S2).

To select the most interesting modules for the aging process, we followed a two-step approach. First, for each module, the corresponding gene expression profiles were summarized into a representative - module eigengene (ME; see *Methods - Identification of significantly age-associated modules, hub genes, and DEG-module-hub overlapping genes*) - and this illustrative profile was correlated with aging. All modules with a significant (false discovery rate (FDR) < 0.05) and at least moderate (≥ 0.4) correlations between their ME and age were selected. The brain exhibited 4 modules significantly correlated with age (3 positive and 1 negative); the heart and the muscle both showed only 1 module negatively correlated with age; the liver displayed 5 modules negatively correlated with age and 4 modules with positive correlation; and the pancreas did not have any modules significantly correlated with age and was not considered in further analyses (Figure 2A).

**Figure 2.**
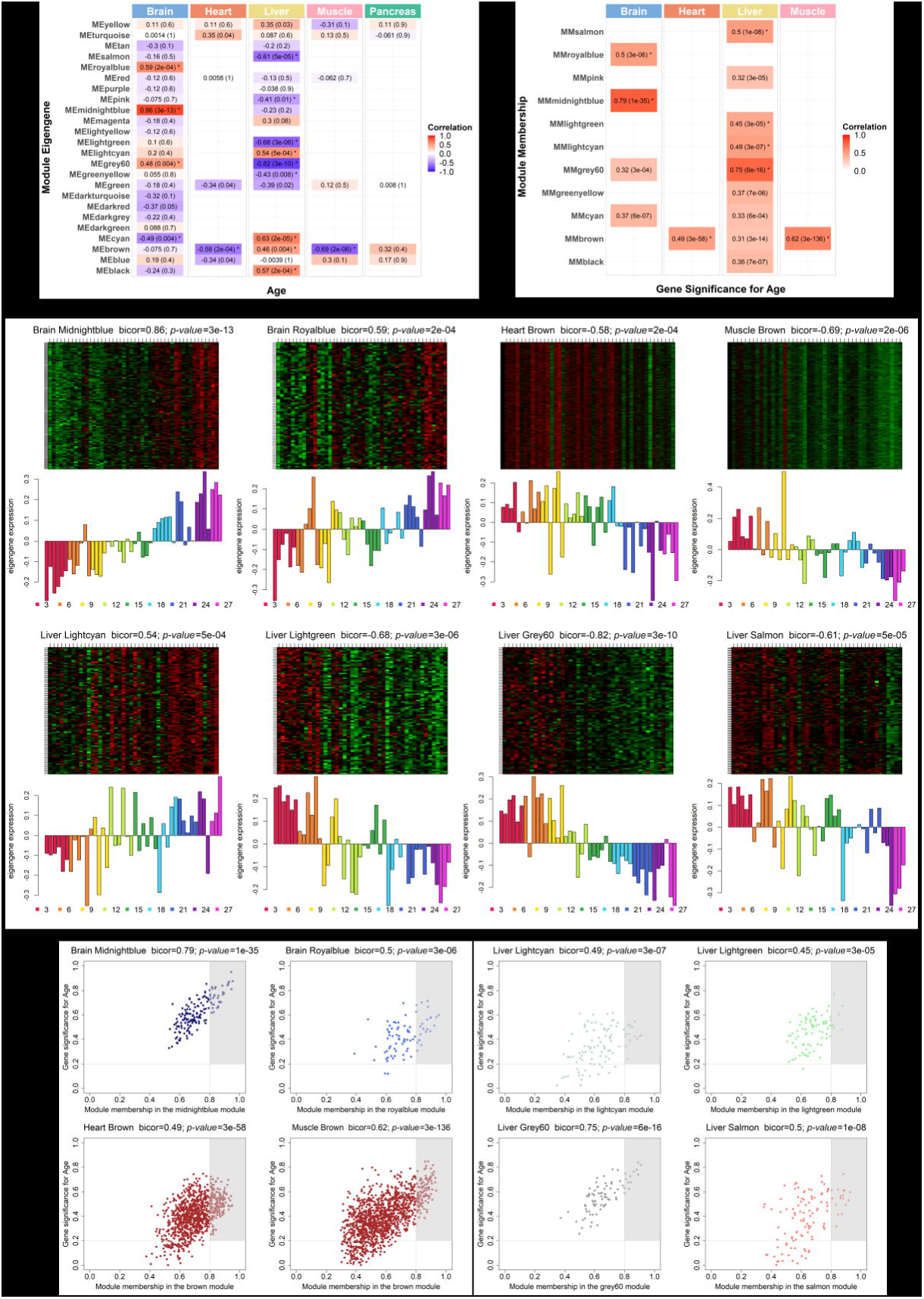
Weighted gene co-expression network analysis of the whole mouse transcriptome. **A**) Correlation between each module’s eigengene (ME) and age. Each tissue exhibits a variable number of modules of co-expressed genes (brain: 24; heart: 6; liver: 18; muscle: 6; pancreas: 5), and unassigned genes are clustered together in the grey module (not shown). ME is the first principal component of the expression matrix of a module, thus being the most representative gene expression profile of that group of correlated genes. Cells are annotated with bicor values and corresponding FDR adjusted *p-values* (inside brackets). Red and blue cells depict positive and negative correlations, respectively. The intensity of color represents the degree of correlation. All modules whose ME’s correlation with age is significantly equal or higher than 0.4 were considered (moderate correlation and above; FDR < 0.05; marked with *). **B**) Correlation between module membership (MM) and gene significance (GS) of the previously selected modules. MM is obtained by correlating the expression of individual genes to the ME, and GS corresponds to the absolute value of the correlation between individual genes and the trait of interest. Only modules with moderate or higher (≥ 0.4) and significant (*p-value* < 0.05) correlations were considered for subsequent analysis (marked with *). **C**) Gene expression profile of each module. The heatmaps (top) display the standardized expression (*z-score*) of individual genes (rows) per sample (columns), whereas the bar plots (below) represent the ME expression profile. Each bar of the bar plot corresponds to the same samples of the heatmap. Negative (positive) values of ME expression relate to the under-expression (over-expression) of genes in each module’s heatmap (green and red colors, respectively). **D**) Intramodular hub gene identification. For each module, genes with individual GS > 0.2 and MM > 0.8 were considered to be the most functionally important (inside grey rectangles).

Next, the gene significance (GS) and module membership (MM) measures were analyzed. GS refers to the absolute value of individual correlations of genes to the trait of interest, whereas MM relates to the individual correlations of genes to the ME. High correlations between these two measures are indicative of genes that are highly significant for aging being as well highly important to the module. Modules exhibiting significant (*p-value* < 0.05) and at least moderate MM-GS correlations (≥ 0.4) were selected. From the 4 previously selected modules in the brain, only 2 exhibited significant MM-GS correlations (*Midnightblue* and *Royalblue* modules); in the heart and muscle, the single selected modules also displayed significant MM-GS correlations (*Brown* module in both tissues); and in the liver, from the 9 pre-selected modules, 4 passed the MM-GS correlation criteria (*Salmon*, *Lightgreen*, *Lightcyan* and *Grey60* modules) (Figure 2B; Supplemental File S2). These selected clusters of genes can be considered the most relevant to the aging process.

In order to evaluate each module’s age-related gene expression, we plotted a heatmap of VST-normalized expression values of the genes belonging to the module, accompanied by a bar plot representing the ME expression profile. This visualization allowed us to better understand the behavior of the age-related genes, as well as to identify the lifespan periods where shifts in expression occur. The brain showed an increasing trend in gene expression with age in both modules, the difference being the onset of expression change. In the *Midnightblue* module (bicor = 0.86; *p-value* = 3*e*-13), the shift from under- to over-expression occurs around the transition from middle- to old-age (15 to 18 months), while in the *Royalblue* module (bicor = 0.59; *p-value* = 3*e*-04) this transition is less defined, but probably occurring later in life, within old-age (18 to 21 months) (Figure 2C, *upper left panels*). The heart and the muscle both exhibited decreasing trends in expression with increasing age; however, the shift points take place in different periods. In the heart (bicor = -0.58; *p-value* = 2*e*-04), the transition from over- to under expression happened at old-age (18 to 21 months), while in the muscle (bicor = -0.69; *p-value* = 2*e*-06), it happened earlier in life, at middle-age (9 to 12 months) (Figure 2C, *upper right panels*). In the liver, 3 modules exhibited decreasing trends in expression throughout life (*Lightgreen*, *Grey60* and *Salmon*), whereas only one has an increasing trend (*Lightcyan*). In the liver’s *Lightcyan* module (bicor = 0.54; *p-value* = 5*e*-04), gene expression starts to increase around the early middle-life (9 to 12 months); while in the *Lightgreen* (bicor = -0.68; *p-value* = 3*e*-06), *Grey60* (bicor = -0.82; *p-value* = 3*e*-10) and *Salmon* (bicor = -0.61; *p- value* = 5*e*-05) modules, gene expression starts to decrease in the transition from mature adulthood to middle-life (6 to 9 months), within middle-age (12 to 15 months) and in the transition from middle- to old-age (15 to 18 months), respectively (Figure 2C, *lower panels*). Furthermore, in each module, we identified the genes with the highest MM and GS – Hub genes - as they are important elements of the module, as well as the most significantly associated with the trait (see *Methods - Identification of significantly age-associated modules, hub genes, and DEG-module-hub overlapping genes*; Figure 2D and Supplemental File S2).

### Altered genes and biological networks provide tissue-specific markers of aging

In order to integrate the results from the two described approaches for finding age-dysregulated genes, we intersected the resulting gene lists and evaluated their functional implications. For each tissue, we first aggregated the DEGs from each pairwise comparison into a single gene set comprising all genes found to be differentially expressed in each time point against the 3-month reference. We then compared this gene set with the other gene sets of interest (TDEGs, Module genes, and Hub genes) and selected for functional analysis the intersection of, at least, the DEGs and the Hub genes (Figure 3; Supplemental File S3). As expected, all TDEGs overlapped with the DEGs, and all Hub genes overlapped with the Module genes.

**Figure 3.**
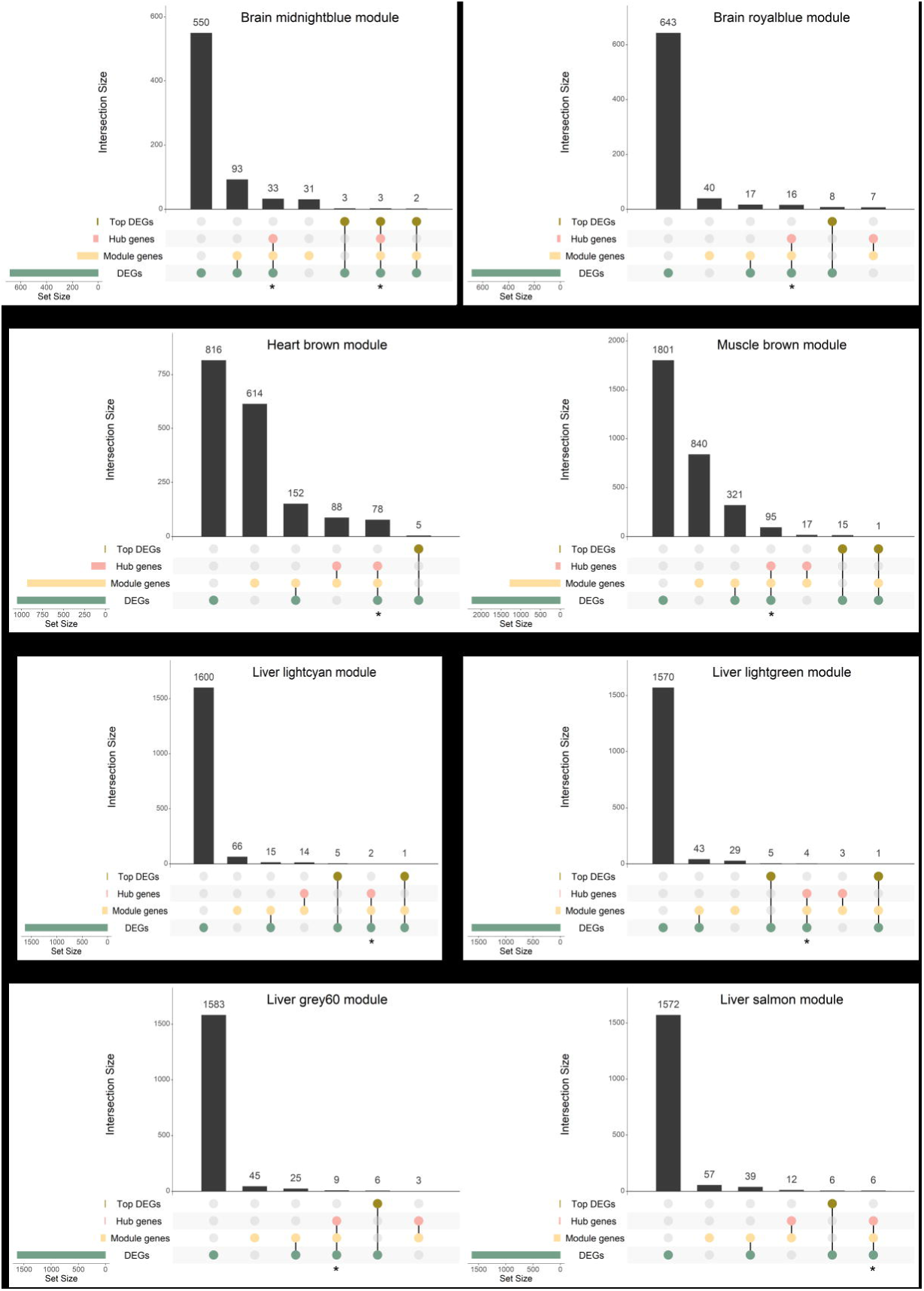
Gene overlap between DEGs, TDEGs, module genes, and hub genes. Bars represent intersection size and colored circles depict the gene sets involved. Genes in common at least in the DEG and hub gene sets were considered for further analysis (identified with *).

In the brain *Midnightblue* module, 36 genes were selected for further analyses, from which 33 resulted from the *DEGs-Module-Hub* intersection, and 3 resulted from the same intersection with an additional layer of TDEGs (Figure 3; Supplemental File S3). As for the rest of the modules, the selected genes resulted exclusively from the *DEGs-Module-Hub* intersection: 16, 78, 95, 2, 4, 9, and 6 genes from the brain *Royalblue*, heart *Brown*, muscle *Brown*, and liver *Lightcyan*, *Lightgreen*, *Grey60* and *Salmon* modules, respectively (Figure 3; Supplemental File S3). All overlapping genes present the same direction of dysregulation in both approaches (Supplemental File S3).

To better understand the functions underlying these signature gene lists, we performed an enrichment analysis on GO BPs and selected the ones with an FDR adjusted p-value of less than 0.05 (see *Methods - Functional characterization of DEG-module-hub genes’ overlap*; Supplemental Figure S2 and Supplemental File S4). To deal with the large number of significant BPs exhibited by some modules and to provide a clear picture of how they and their associated genes relate to each other, we constructed a network of these results, with nodes representing GO terms and edges depicting gene overlap (see *Methods - Network visualization of functionally enriched terms*). After having constructed the network, we addressed GO term redundancy by clustering together nodes based on gene overlap similarity and then assigning automatically created labels from the most frequent words in the cluster, as well as words adjacent to the most represented ones (see *Methods - Network visualization of functionally enriched terms*; Supplemental File S4). To further simplify the visualization, we created a summary network based on the generated clusters, where all the nodes belonging to the same cluster collapsed into a *meta-node* and all the edges connecting the different clusters collapsed into *meta-edges* as well [43] (Supplemental Figure S2; Supplemental File S4). Selected results are present in Table 2.

**Table 2.**
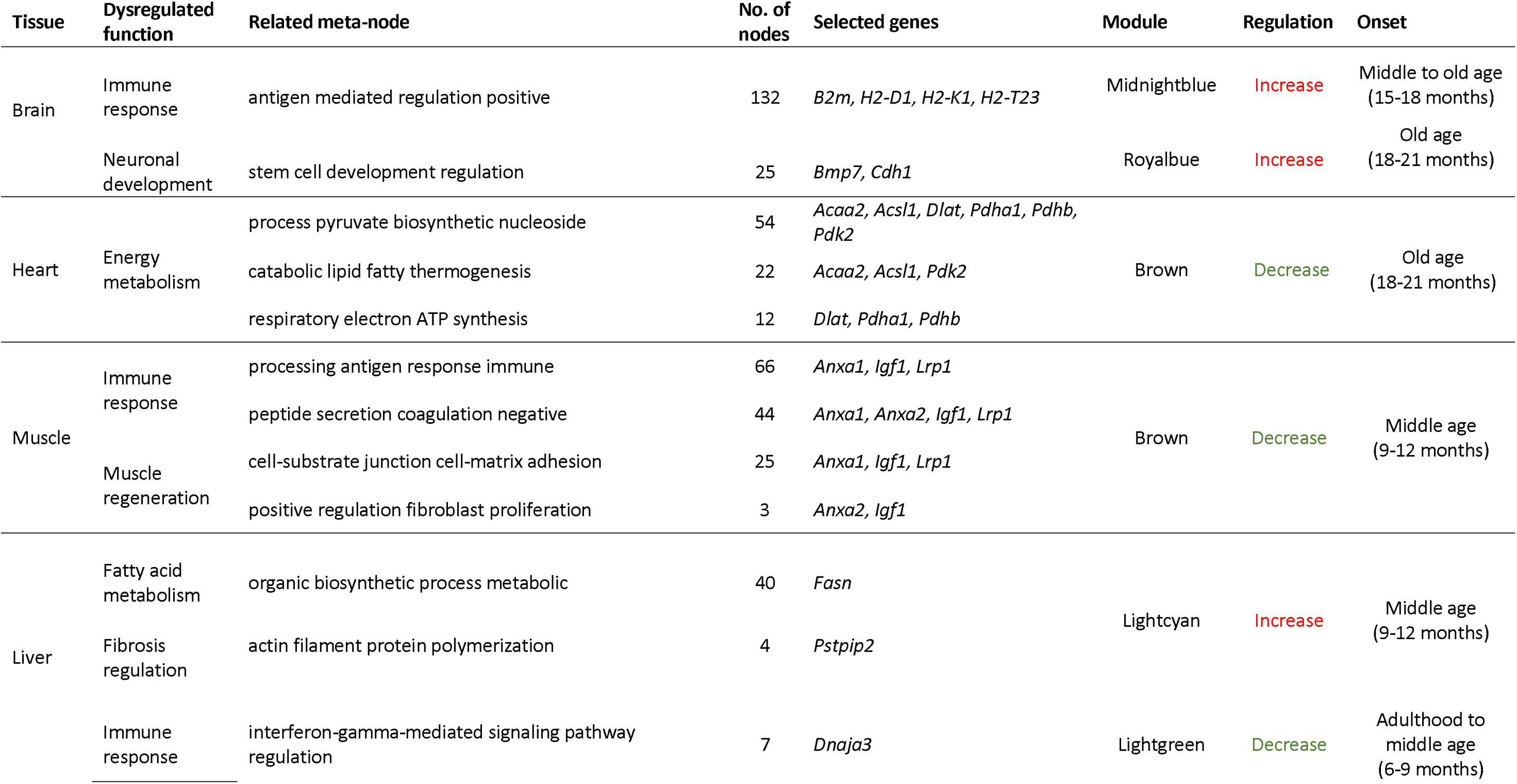
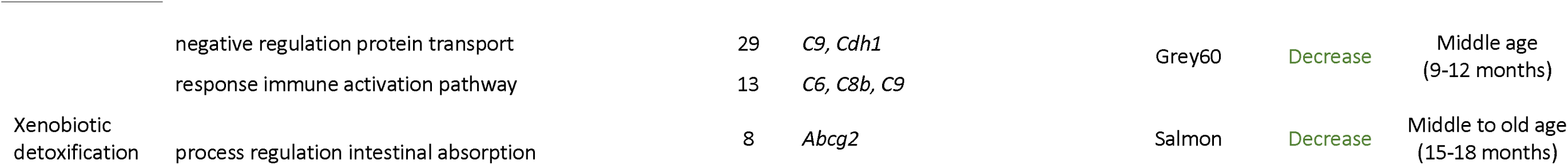
Tissue-specific age-dysregulated functions. Relates to Supplemental Figure S2 and Supplemental File S4.

#### Immune response processes and stem cell development are upregulated in the aging brain

GO enrichment analysis of the overlap of the brain DEGs with the *Midnightblue* module genes identified 224 significantly over-represented BPs, allocated into 9 *meta-nodes* (Supplemental Figure S2A – Brain *Midnightblue* module; Supplemental File S4). The largest *meta-node* is involved in antigen-mediated immunity and comprises 132 GO terms, thus representing more than half of all significant processes observed for this set (Table 2 – Brain *Midnightblue* module; Supplemental File S4). The genes present in this group exhibit increased expression with increasing age, shifting from down- to up-regulation around the transition from middle to old age (15-18 months; Figure 2C – Brain *Midnightblue* module; Table 2 – Brain *Midnightblue* module). Among the genes identified within this cluster, several encode for Major Histocompatibility Complex I (MHCI) proteins, such as in the case of *H2-T23* (*H-2 class I histocompatibility antigen D-37 alpha chain*), *H2-D1* (*histocompatibility 2, D region locus 1*), *H2-K1* (*histocompatibility 2, K1, K region*) and *B2m* (*beta 2 microglobuli*n) (Table 2 – Brain *Midnightblue* module; Supplemental File S4).

Furthermore, 25 processes related to stem cell development were also found to be significantly enriched in the aging brain, this time regarding the gene set resulting from the intersection of brain DEGs and the *Royalblue* module genes (Table 2 – Brain *Royalblue* module; Supplemental File S4). This *meta-node* (Supplemental Figure S2A – Brain *Royalblue* module) includes genes whose expression also tend to increase throughout aging but the shift from down- to up-regulation occurs later in life, within old age (18-21 months; Figure 2C – Brain *Royalblue* module; Table 2 – Brain *Royalblue* module). Among these clusters’ genes, we highlight *Bmp7* (*bone morphogenetic protein 7*), and *Cdh1* (*cadherin 1*) (Table 2- Brain *Royalblue* module; Supplemental File S4).

#### Decline of cardiac energy metabolism with age

In the heart, we identified 12 clusters of similar GO terms (Supplemental Figure S2B – Heart *Brown* module), comprising 133 significantly enriched GO BPs based on the intersection between the heart DEGs and the heart *Brown* module genes, from which approximately two-thirds (n = 88) are directly related to energy production (Table 2 – Heart *Brown* module; Supplemental File S4). The genes found to be involved in these processes exhibit decreased expression along the lifespan, with the transition from up- to down-regulation occurring late in life (18-21 months; Figure 2C – Heart *Brown* module; Table 2 – Heart *Brown* module). These genes include *Pdha1* and *Pdhb* (*pyruvate dehydrogenase E1 alpha 1* and *pyruvate dehydrogenase (lipoamide) beta*, respectively), *Dlat* (*dihydrolipoamide S-acetyltransferase (E2 component of pyruvate dehydrogenase complex)*), *Pdk2* (*pyruvate dehydrogenase kinase, isoenzyme 2*), *Acsl1* (*acyl-CoA synthetase long-chain family member 1*), and *Acaa2* (*acetyl-Coenzyme A acyltransferase 2 (mitochondrial 3-oxoacyl-Coenzyme A thiolase)*) (Table 2 – Heart *Brown* module; Supplemental File S4).

#### Immune and regeneration processes are altered in the muscle of aged mice

Over-representation analysis of GO terms in the gene set obtained from the overlap of the muscle DEGs and the muscle *Brown* module genes resulted in a highly interconnected network of 334 BPs organized into 22 *meta-nodes* (Supplemental Figure S2C – Muscle *Brown* module; Supplemental File S4). The largest cluster comprises 66 GO terms and includes genes related to antigen processing and immunity (Table 2 –Muscle *Brown* module; Supplemental File S4). Moreover, muscle regeneration processes were also enriched in the aging muscle (Table 2 – Muscle *Brown* module: Supplemental File S4). The genes present in this set also exhibit decreased expression with increasing age, shifting from up- to down-regulation relatively early in the lifespan, within middle age (9-12 months; Figure 2C – Muscle *Brown* module; Table 2 – Muscle *Brown* module). Among these genes, we highlight *Igf1* (*insulin-like growth factor 1*), *Anxa1* and *Anxa2* (*Annexin A1* and *A2*), and *Lrp1* (*low density lipoprotein receptor-related protein 1*), highly relevant in the context of aging and longevity studies (Table 2 – Muscle *Brown* module; Supplemental File S4).

#### Liver aging is characterized by a global dysregulation of hepatic function

In the liver, a total of 113 BPs were identified as being significantly enriched among the gene lists resulting from the overlap of the liver DEGs and module genes (Table 2; Supplemental File S4). In the DEG-*Lightcyan* module gene set, 44 enriched GO terms were allocated into 2 *meta- nodes*, with the largest (n = 40) relating to fatty acid metabolism and the smallest (n = 4) to hepatic fibrosis regulation (Supplemental Figure S2D – Liver *Lightcyan* module; Table 2 – Liver *Ligthcyan* module; Supplemental File S4). The genes involved in these processes – *Fasn* (*fatty acid synthetase*) and *Pstpip2* (*proline-serine-threonine phosphatase-interacting protein 2*) – exhibit increased expression throughout the lifespan, with the shift from down- to up- regulation occurring within middle age (9-12 months; Figure 2C – Liver *Lightcyan* module; Table 2 – Liver *Lightcyan* module). Regarding the DEG-*Lightgreen* module gene overlap, 2 clusters of processes were identified related to immune function, among which we highlight one *meta-node* comprising 7 interferon-gamma protein signaling-related GO terms (Supplemental Figure S2D – Liver *Lightgreen* module; Table 2 – Liver *Lightgreen* module; Supplemental File S4). The identified genes within these clusters include *Dnaja3* (*DnaJ heat shock protein family (Hsp40)*) and present a decreased trend in expression with increasing age, shifting from up- to down-regulation around the transition from mature adulthood to middle age (6-9 months; Figure 2C – Liver *Lightgreen* module; Table 2 – Liver *Lightgreen* module). Furthermore, GO term enrichment analysis in the gene list obtained from the intersection of the liver DEGs and the *Grey60* module genes resulted in 2 *meta-nodes*, comprising processes regarding the activation of immune responses (Supplemental Figure S2D – Liver *Grey60* module; Table 2 – Liver *Grey60* module; Supplemental File S4). The genes involved in these processes include *C6*, *C8b*, *C9* (*complement components 6, 8 beta polypeptide,* and *9*), and *Cdh1* and also exhibit decreased expression throughout the lifespan, with the shift from up- to down-regulation occurring within middle age (9-12 months; Figure 2C – Liver *Grey60* module; Table 2 – Liver *Grey60* module). Finally, in the DEG-*Salmon* module gene overlap, we identified 18 significantly enriched GO BPs, allocated into 3 clusters (Supplemental Figure S2D – Liver *Salmon* module). Among these clusters, we highlight the one related to xenobiotic detoxification (n = 8) (Table 2 – Liver *Salmon* module; Supplemental File S4) The gene involved in these processes – *Abcg2* (*ATP binding cassette subfamily G member 2 (Junior blood group)*) - exhibits decreased expression across aging, transitioning from up- to down-regulation in the transition from middle to old age (15-18 months; Figure 2C – Liver *Salmon* module; Table 2 – Liver *Grey60* module).

### Similar biological processes are altered with aging among tissues

So far, the evidence pointed to a greater contribution of tissue type, rather than age, to gene expression variation between the samples and, for that reason, all the analyses were performed in each tissue independently. In fact, very few genes that were found to be key players of aging are shared between tissues, in a maximum of two organs simultaneously (Brain:Muscle - 8, Brain:Liver - 1, Heart:Liver - 1; Figure 4 - right panel; Supplemental File S3). Nonetheless, when we compared the significantly enriched GO terms between each tissue, we found a much higher overlap than that observed at the gene level (Figure 4 - left panel; Supplemental File S5). The highest number of shared BPs (65) was observed between the brain and the muscle, followed by 19 processes shared between the heart and the liver. The brain, the muscle and the liver exhibited 12 GO terms in common, while the muscle and the liver shared 7 BPs, and the brain and the liver displayed 6 processes in common. Finally, the brain, the heart and the liver, presented only 1 BP in common, as was the case of the muscle and the heart (Figure 4 – left panel; Supplemental File S5). As described above, we organized the obtained GO terms in summary networks (Supplemental Figure S3; Supplemental File S5) and organized the selected results into a table (Table 3).

**Figure 4.**
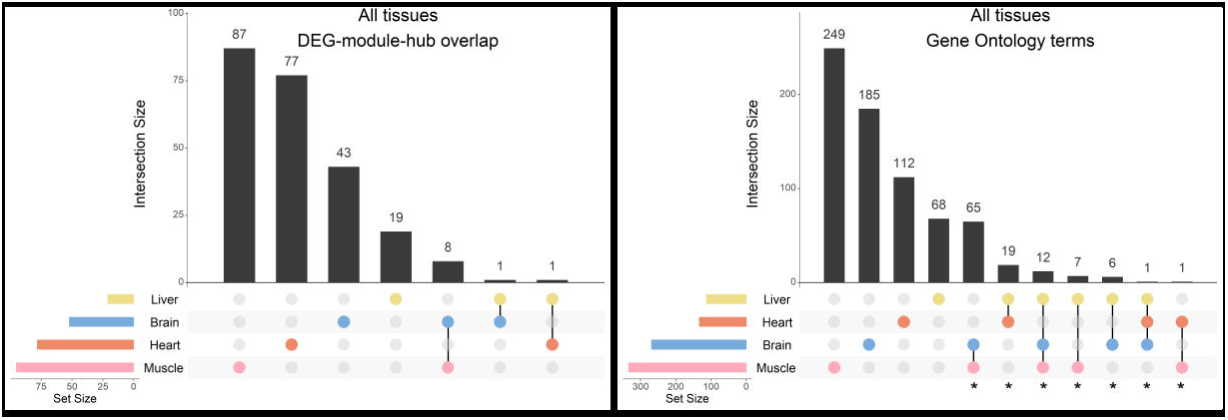
Overlap of genes and biological processes between the brain, heart, liver, and muscle. Upset plots depicting the gene overlap between DEGs, module genes, and hub genes per tissue (left), as well as the overlap of the enriched GO terms in the same tissues (rigth). Bars represent intersection size and colored circles depict the gene/GO term sets involved. Each tissue’s gene list results from the intersection of DEGs, module, and hub genes. In tissues with more than one module (i.e. the brain and the liver), the gene list results from the combination of each module’s intersection, and the GO term list results from the combination of each module’s GO terms. GO terms in common at least in two tissues were considered for further analysis (identified with *).

**Table 3.**
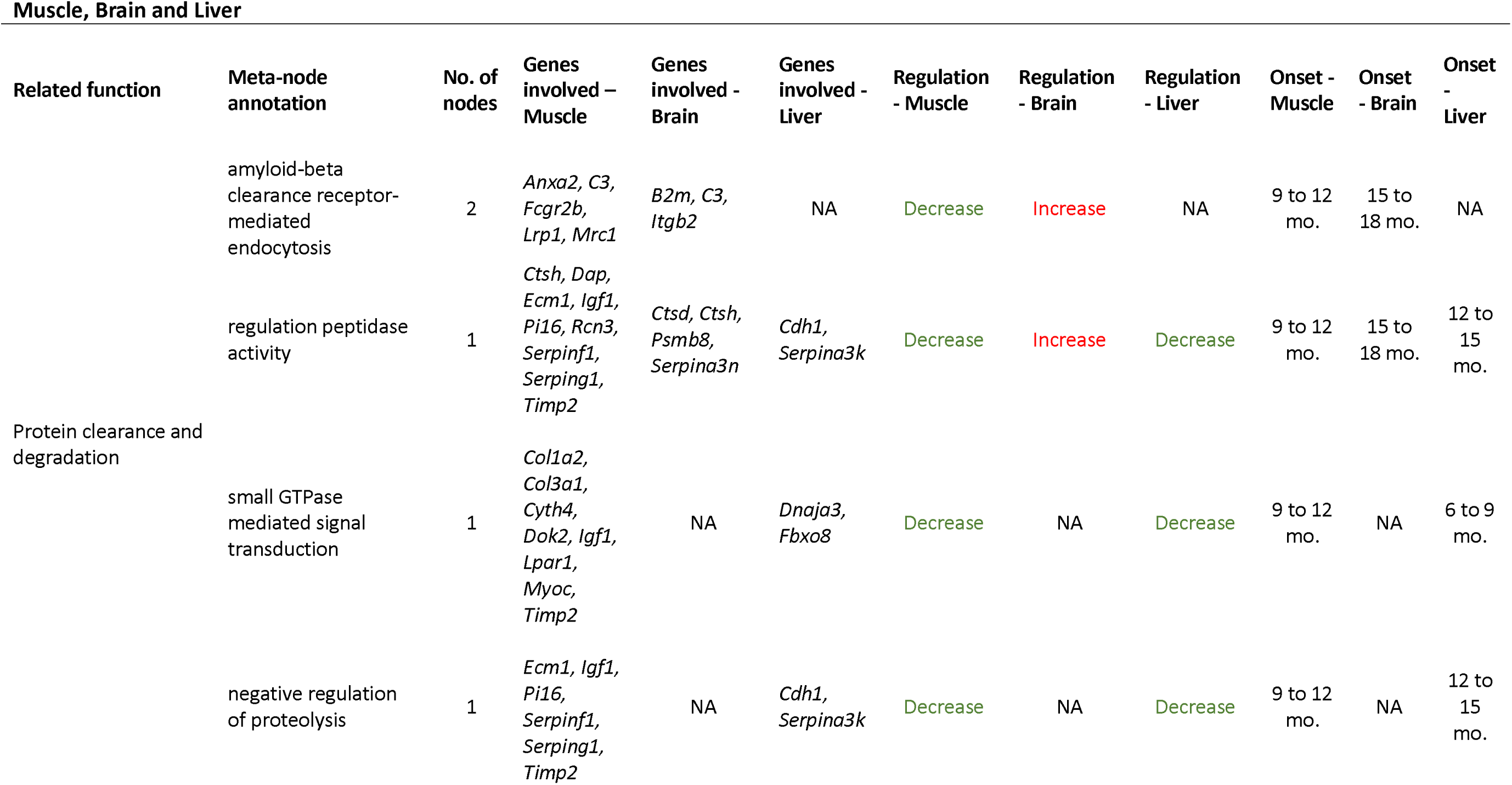
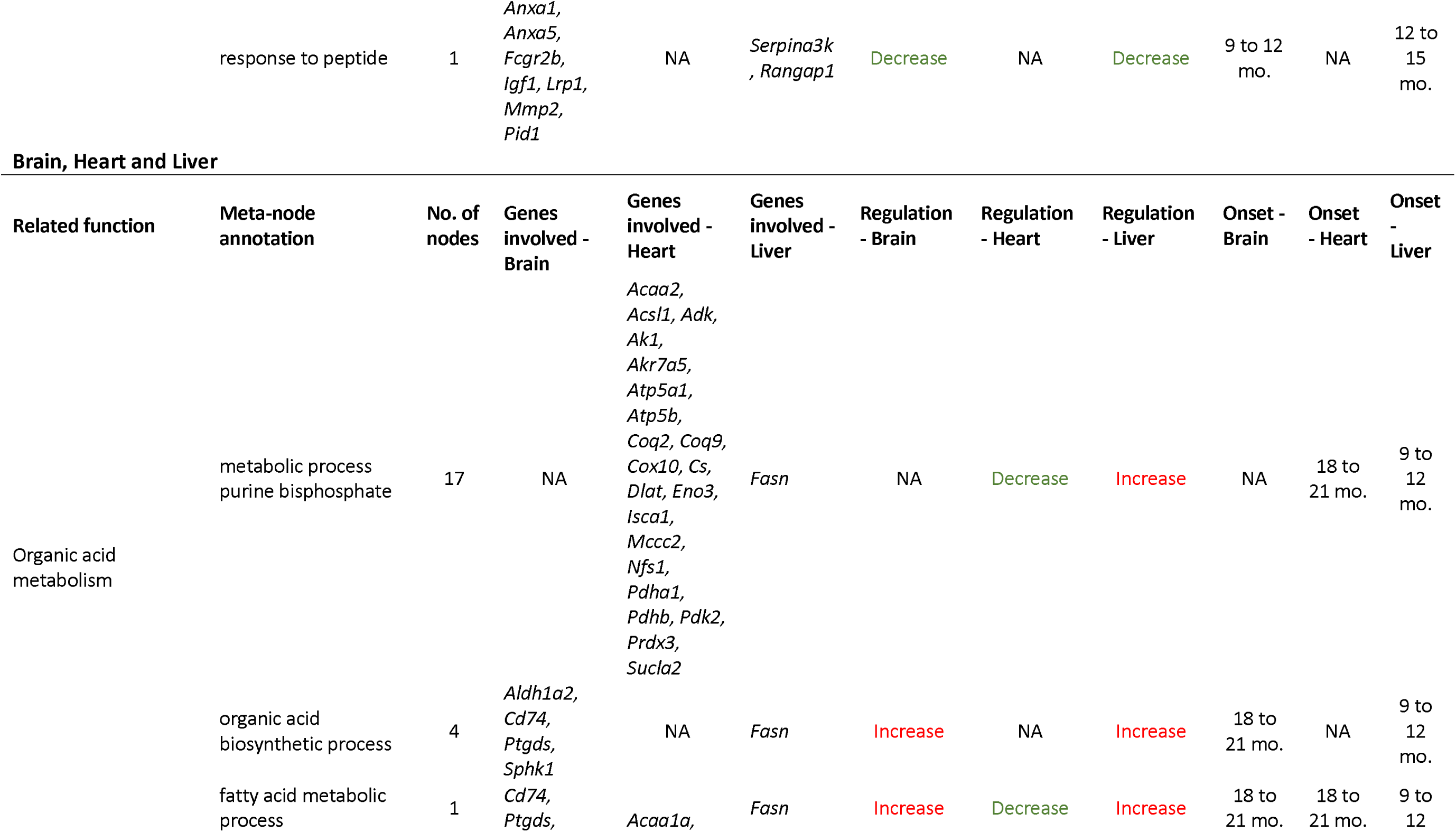
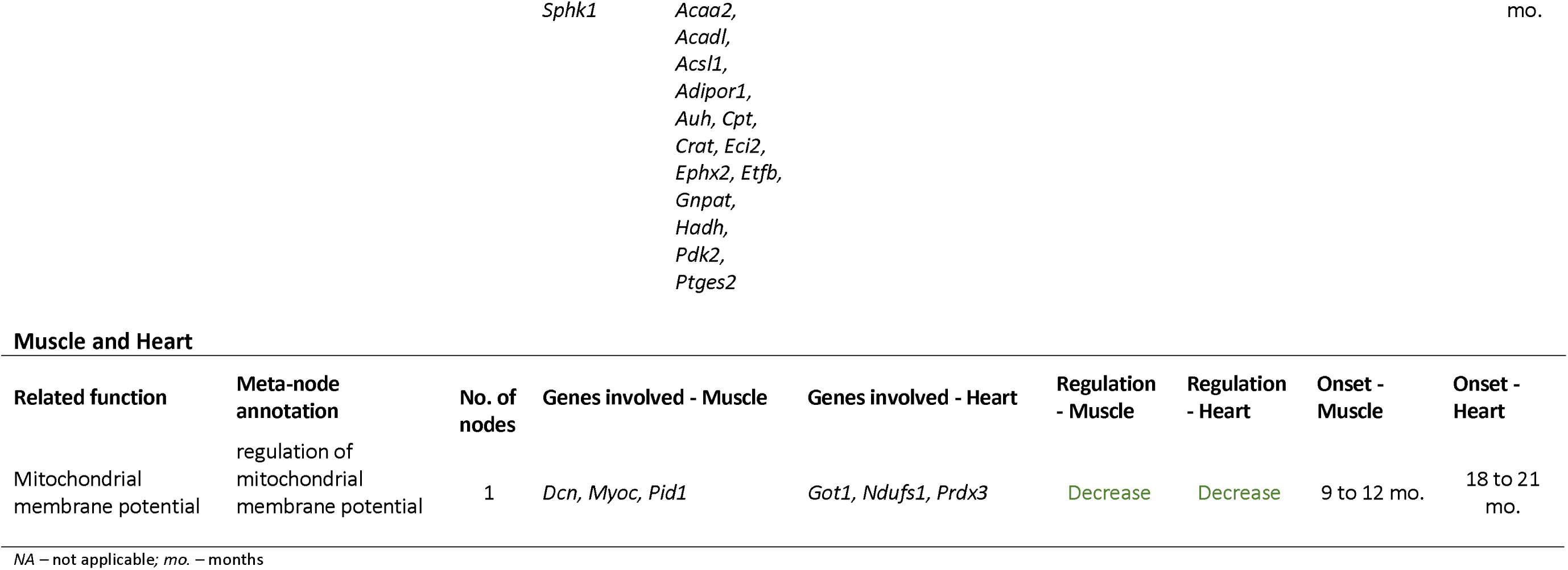
Inter-tissue age-dysregulated functions. Relates to Supplemental Figure S3 and Supplemental File S5.

#### Protein clearance and degradation processes are compromised in the aging brain, muscle and liver

The muscle and the brain share 65 GO biological processes clustered into 12 *meta-nodes* (Supplemental Figure S3 – Muscle:Brain; Supplemental File S5). In line with what we observed at the tissue-level, immune processes are overly represented in the intersection of GO terms between these two tissues (38 out of 65 BPs, approx.58%; Supplemental File S5). Among the other identified shared processes, we highlight the clearance of Amyloid-β (Aβ) by receptor-mediated endocytosis (Table 3 – Muscle:Brain; Supplemental File S5). In the muscle, the genes involved in the two processes comprising this *meta-*node exhibit a decreasing trend of expression across the lifespan, transitioning from up- to down-regulation within middle age (9- 12 months; Figure 2C – Muscle *Brown* module; Table 3 – Muscle:Brain). Conversely, in the brain, the genes implicated in these processes are increasingly expressed with aging, with the shift from down- to up-regulation occurring slightly later in life, marking the transition from middle- to old-age (15-18 months; Figure 2C – Brain *Midnightblue* module; Table 3 – Muscle:Brain).

In the network representing the overlapping processes between the muscle and the liver, each of the 7 shared BPs constituted a single *meta-node* (Supplemental Figure S3 – Muscle:Liver; Supplemental File S5). Interestingly, 3 out of the 7 BPs are also related to proteostasis (Table 3 – Muscle:Liver; Supplemental File S5), an observation in line with the findings from the Muscle:Brain overlap. Furthermore, the genes implicated in these processes exhibit decreased expression in both tissues, with the shift from up- to down-regulation occurring around or within middle age (6-9 and 12-15 months in the liver and 9-12 months in the muscle; Figure 2C – Muscle *Brown* and Liver *Lightgreen* and *Grey60* modules; Table 3 – Muscle:Liver). The muscle, the brain and the liver exhibit 12 shared GO terms grouped together in three different *meta-nodes* (Supplemental Figure S3 – Muscle:Brain:Liver; Supplemental File S5). Similarly to the Muscle:Brain overlap, immunity-related pathways are largely over-represented in this overlap as well, with their associated *meta-node* comprising 83% of all shared BPs (10 out of 12 BPs; Supplemental File S5). Among the other two clusters of BPs, we highlight the regulation of peptidase activity (Table 3 – Muscle:Brain:Liver; Supplemental File S5), as it is in line with the Muscle:Brain and Muscle:Liver results. The brain is the only tissue exhibiting increased expression throughout life of genes involved in this process, however the expression shift occurs the latest, in the transition from middle- to old age (15-18 months; Figure 2C – Brain *Midnightblue* module; Table 3 – Muscle:Brain:Liver). Both the muscle and the liver exhibit decreased gene expression associated with this process, shifting within middle age (9- 12 and 12-15 months, respectively; Figure 2C – Muscle *Brown* and Liver *Grey60* modules; Table 3 – Muscle:Brain:Liver).

#### Dysregulation of organic acid metabolism across the lifespan is shared by the brain, the heart and the liver

Unsurprisingly, the commonly affected BPs with aging in the heart and the liver are related to metabolism, with the 19 identified shared processes allocated into 2 clusters related to purine and glycerol ether metabolism (Supplemental Figure S3 – Heart:Liver; Supplemental File S5). Notably, the GO terms related to purine metabolism represent 89% of all shared BPs (17 out of 19 BPs; Table 3 – Heart:Liver; Supplemental File S5) and the genes comprised by this *meta-node* in the heart display decreased expression with increasing age, whereas the liver genes involved in the same processes exhibit increased expression (Figure 2C – Heart *Brown* and Liver *Lightcyan* modules; Table 3 – Heart:Liver). Moreover, the shifts in regulation occur late in life in the heart, within old age, whereas in the liver they are observed earlier, within middle age (18-21 and 9-12 months, respectively; Figure 2C – Heart *Brown* and Liver *Lightcyan* modules; Table 3 – Heart:Liver). Intersecting biological processes identified in the liver and the brain resulted in 6 shared GO terms, clustered into 3 *meta-nodes* (Supplemental Figure S3 – Brain:Liver), with the largest cluster comprising approximately 67% of all BPs (4 out of 6) and relating to the biosynthesis of organic acids (Table 3 – Brain:Liver; Supplemental File S5). In both tissues, the genes involved in these processes are increasingly expressed with aging, however shifting from down- to up-regulation at different lifespan time points (18-21 and 9-12 months in the brain and the liver, respectively; Figure 2C – Brain *Royalblue* and Liver *Lightcyan* modules; Table 3 – Brain:Liver). The overlap of processes between the brain, the heart and liver consists of a single BP and *meta-node* related to fatty acid metabolism (Supplemental Figure S3 – Brain:Heart:Liver; Table 3 – Brain:Heart:Liver; Supplemental File S5). The genes involved in this process display an increasing trend of expression throughout the lifespan in both the brain and the liver, however with very different onsets of expression change (9-12 and 18-21 months in the liver and brain, respectively; Figure 2C – Brain *Royalblue* and Liver *Lightcyan* modules; Table 3 – Brain:Heart:Liver). Conversely, the genes involved in this process in the heart have decreased expression, transitioning from up- to down-regulation within old-age (18-21 months; Figure 2C – Heart *Brown* module; Table 3 – Brain:Heart:Liver).

#### Age affects the expression of genes involved in mitochondrial membrane potential in the heart and the muscle

The single age-related dysregulated process in common between the muscle and the heart is related to mitochondrial activity, particularly to the regulation of membrane potential (Supplemental Figure S3 – Muscle:Heart; Table 3 – Muscle:Heart; Supplemental File S5). Notably, even though both tissues exhibit the same decreasing trend in the expression of genes related to this process, in the muscle the transition from up- to down-regulation occurs much earlier in life than in the heart (9-12 and 18-21 months, respectively; Figure 2C – Muscle *Brown* and Heart Brown modules; Table 3 – Muscle:Heart).

## Discussion

In this work, we provide an in-depth characterization of the age-associated alterations in gene expression throughout the murine lifespan. More specifically, we employed two approaches to identify genes central to the aging process with a higher degree of certainty. First, we performed differential gene expression profiling between each lifespan time point (6, 9, 12, 15, 18, 21, 24 and 27 months) relative to the reference time point (3 months) and selected the DEGs in each pairwise comparison. Then, the network-based approach allowed us to establish tissue-specific clusters of co-expressed genes that significantly correlated with increasing age. By integrating the results of these two methodologies, we were able to establish gene expression signatures of aging with greater confidence, and because we did not apply any fold-change threshold in the selection of DEGs, these signatures include genes whose expression change is small but potentially relevant at the proteome level.

Among the selected tissues, we found that there are modest tissue-specific changes in gene expression, and of those that we studied, the pancreas was the only tissue that appears to be largely unaffected by aging, a finding that was corroborated both by the low yield of DEGs and by the absence of significantly age-associated gene co-expression modules. Unsurprisingly, the observed gene expression alterations across the lifespan generally reflect loss of tissue function and homeostasis (Supplemental Discussion). However, lipid transport and metabolism may also play an important role in organismal aging with at least one TDEG related to these processes being identified in 4 of the 5 studied tissues (Supplemental Discussion). Moreover, different tissues exhibit diverse onsets of gene expression dysregulation, thus evincing an asynchronous age-related dysregulation of gene expression, in line with recent findings [6, 20].

Although very little commonalities in age-related dysregulation were observed at the gene-level, we realized that the studied tissues could be grouped accordingly by shared biological processes affected by aging. The brain, heart and liver share age-related dysregulation of processes related to the metabolism of organic acids, namely fatty acids, while the heart and muscle share dysregulation of mitochondrial membrane potential (Supplemental Discussion).

Interestingly, the aging brain, muscle and liver share dysregulation of processes related to protein clearance and degradation mechanisms, among others. It is well known that proteostasis decline is a hallmark of aging [44], leading to proteome imbalances and contributing to protein aggregation, including amyloid-like aggregates, in both muscle and the brain [45–48]. As a result, age-related protein aggregates also increase the risk for Aβ aggregation, a pathological hallmark in Alzheimeŕs disease (AD) [45–48]. In our observations, age-related dysregulation in the clearance of Aβ by receptor-mediated endocytosis is shared by the brain and the muscle. In the brain, *complement component 3* (*C3*) has been reported to mediate the phagocytosis and clearance of insoluble Aβ by microglial cells in C57BL/6 mice [49]. However, another study reported decreased levels of Aβ internalization in microglia from aged mice when compared to younger individuals [50]. Taken together, our observations of increased expression of *C3* during physiological aging in the brain may reflect an initial compensatory increase in phagocytic clearance of Aβ in response to the physiological accumulation of these peptides, most likely followed by a decrease in this capacity due to the overburden of cytotoxic aggregates.

Notably, Aβ aggregates have been implicated in age-related protein conformational diseases in other tissues, such as the skeletal muscle [51, 52]. Sporadic inclusion-body myositis (s-IBM) is a degenerative muscle disease for which age is a major risk factor and it features progressive muscle-fiber degeneration which is characterized by the accumulation of multiple misfolded protein aggregates including Aβ peptides [reviewed in 52]. We found the expression of some genes involved in Aβ clearance by receptor-mediated endocytosis to decrease around mid-life in this tissue, including that of *C3*, *Fcgr2b* (*Fc receptor, IgG, low affinity IIb*) and *Lrp1*. These genes have already been described as important players of Aβ internalization in the brain by glial cells and macrophages [53, 54], which suggests that the accumulation of Aβ in old age in skeletal muscle may be preceded by the middle-age down-regulation of Aβ internalization most likely by resident macrophages.

Moreover, both proteasomal and autophagy–lysosomal pathways may also be particularly dysregulated with age in these tissues, as we observed increased expression of *Psmb8* (*proteasome (prosome, macropain) subunit, beta type 8 (large multifunctional peptidase 7)*) and *Ctsd* (*cathepsin D*) in the brain, and decreased expression of *Igf1, Ctsh* (*cathepsin H*), and *Dap* (*death-associated protein*) in the muscle, and of *Dnaja3* and *Fbxo8* (*F-box protein 8*) in the liver. In the brain, the proteasome subunit *Psmb8* and the lysosomal protease *Ctsd* are both involved in maintaining proteostasis, as essential players in proteasomal and autophagic activities, respectively [55, 56]. *Psmb8* is critical for immunoproteasome assembly and is upregulated in long-lived primate species and human fibroblasts [55]. The immunoproteasome is also involved in protein degradation, especially in neurodegenerative diseases such as Lewy Body Dementia (LBD) and AD, where there is also elevated expression of *Psmb8* [57, 58]. Heightened proteasomal subunit expression, as we observe with *Psmb8* in the brain, can lead to the formation of dysfunctional proteasome complexes with altered activity, stimulating a compensatory response to cope with defective proteasome function and altered proteostasis [58]. In line with the results seen in long-lived primate species, augmented expression of proteasome subunits has also been shown to extend lifespan in worms and yeast [59]. Interestingly, the observed trajectory of expression of *Psmb8* and *Ctsd* during brain aging, shifting from down- to upregulation around the transition from middle to old age, indicates that immunoproteasome and autophagy may be active in later stages of the mouse lifespan in this tissue.

Moreover, in the muscle, Igf1 inhibition has been shown to increase lysosomal proteolysis in this tissue [60, 61], which, together with our observation of decreased expression of this gene across the murine lifespan, may indicate that lysosomal proteolysis is heightened with age in mice. As for *Ctsh*, despite not being largely studied in the skeletal muscle, its age-associated downregulation reported by us may indicate that cathepsin H-mediated protein degradation also declines with age, as this gene encodes for a lysosomal cysteine protease recently implicated in mediating degradation leading to liver fibrosis [62]. Moreover, DAP1, the protein encoded by the human orthology of *Dap*, was found to negatively regulate autophagy [63], while lower expression of Dap has also been linked with alterations in regulation of apoptosis and autophagy, resulting in poor clinical outcomes in human cancers [64, 65]. In summary, these findings suggest an age-related autophagy dysregulation in the aging muscle.

Lastly, despite the lack of evidence regarding the role of *Dnaja3* in the aging liver, this gene encodes a DNAJ/Hsp40 family member, a chaperone/cochaperone complex known to govern the refolding of newly synthesized as well as damaged proteins, targeting ubiquitinated unfolded and/or misfolded proteins for proteasomal degradation [66]. Additionally, *Fbxo8* encodes for a member of the F-box protein family that are part of the ubiquitin E3 ligase SKP1- cullin-F-box (SCF) complex which recognize and recruit target proteins for ubiquitination and degradation by the proteasome [67]. Notably, we found the decrease in expression of these genes to occur relatively early in the lifespan, starting to be observed in the transition from mature adulthood to middle age, around 30-42.5 human years [33]. These observations suggest an early decline in ubiquitination and degradation of unfolded, misfolded and/or damaged proteins in the aging liver and are in line with previous reports of ER chaperone activity in aged mouse livers [68, 69].

Interestingly, serpin family genes are altered in all three tissues, with the downregulation of *Serpinf1 (serine (or cysteine) peptidase inhibitor, clade F, member 1*) and *Serping1 (serine (or cysteine) peptidase inhibitor, clade F, member 1*) in the muscle and *Serpina3n* (serine (or cysteine) peptidase inhibitor, clade A, member 3N) in the brain contrasting with the upregulation of *Serpina3k* (serine (or cysteine) peptidase inhibitor, clade A, member 3K) in the liver. Serpins are a family of serine (or cysteine) protease inhibitors involved in several biological functions including homeostasis control [70]. Notably, *Serpina3n* has been linked with increased immune response activity, is upregulated in aging astrocytes throughout the brain [71] and has also been implicated in AD [addressed in 72]. In addition, Serpina3n mRNA and protein levels are upregulated in a Prion disease mouse model [73]. In the muscle, both *Serpinf1* and *Serping1* are involved in muscle growth and function through regulation of Akt and FoxO signaling pathways [74–76], while in the liver *Serpina3k* has been described as an inhibitor of tissue kallikrein (TK) proteolytic activity, and a modulator of inflammation in the murine liver [77]. Subsequent reports should look to elucidate the mechanisms involved in the age-related expression alterations of serpin-associated genes in these tissues, particularly in the muscle and liver.

This work opens new research avenues as it highlights many unexplored genes and mechanisms in the context of healthy aging and temporally contextualizes gene expression alterations. We identified tissue-specific key players of aging and addressed the functional implications of their age-related alterations. We further identified groups of tissues based on shared age-affected processes, potentially uncovering tissue axes of common age-related functional dysregulation. We found that proteostasis impairment is a common feature of aging in the brain, muscle, and liver. Because these alterations occur at a transcriptional level and protein abundances can be post-transcriptionally regulated, we are currently working on integrating these results with translatomic and proteomic data to comprehensively understand this phenomenon and successfully promote healthy aging strategies.

## Methods

### Dataset characterization

The mouse bulk RNA-Seq data used in this study was made publicly available by the *Tabula Muris Consortium* [6, 35], and is deposited in NCBI’s Gene Expression Omnibus (GEO) under the GEO Series accession number GSE132040 [36]. The original dataset consists of transcriptomic data from 17 male and female mouse tissues across 10 time points (1, 3, 6, 9, 12, 15, 18, 21, 24, and 27 months). For this study, we excluded the 1-month-old samples to avoid the influence of developmental genes [33], and selected the brain, heart, muscle, liver, and pancreas for further analysis (Supplemental File S6). The RNA extraction, cDNA library preparation, RNA sequencing, read quality control, pre-processing and alignment, transcriptome reconstruction, and expression quantification steps are reported elsewhere [6,36,78].

### Differential gene expression analysis

Samples with library size smaller than 4.000.000 reads across all genes were discarded, as described in Schaum *et al.* [6] (Supplemental File S6). Outlier samples were identified based on the sample network approach [79, 80] and excluded if their standardized connectivities (z.K) were more than 2 standard deviations away from the mean z.K (Supplemental File S6; Supplemental Figure S4).

Gene symbols were associated with Ensembl biotype annotations (release 99) [81] using the R package biomaRt (v. 2.44.0) [82, 83]. Differential expression analysis was carried out using DESeq2 (v. 1.28.1) [84] with ‘Age’ as the variable of interest (3-month time point as reference level) and ‘Sex’ as a co-variable (Supplemental Figure S1). Low count genes were pre-filtered and only genes with total read count higher than 10 were kept. Read count data was normalized and transformed with DESeq2’s *estimateSizeFactors* and *vst* functions [85]. PCAs of all original 17 tissues based on the 500 genes with highest row-wise variance (i.e., across all samples) was performed to identify the highest contributing sources of variance. Since samples segregate mainly by tissue (Figure 1A), all subsequent analyses were conducted separately for the brain, heart, liver, muscle, and pancreas (see *Methods - Dataset characterization*).

DEGs were obtained by comparing every time point against the 3-month reference expression level. The Approximate Posterior Estimation for generalized linear model method (apeglm) [86] was used to estimate shrunken log2 fold-changes (log2FC). Genes with *s-values* [87, 88] smaller than 0.005 were identified as significantly differentially expressed [89]. ‘Top DEGs’ (TDEGs) functional annotation was performed using the AmiGO2 webtool [90] (Supplemental File S1).

### Gene co-expression network construction and module construction

The WGCNA R package (v. 1.69) [91] was used to construct co-expression networks for the VST-normalized expression data (see *Methods – Differential gene expression analysis*). Additionally, in order to parallel the use of sex as a co-variable in the differential expression analysis, the normalized expression values were adjusted for this effect using the *removeBatchEffect* function from the Limma R package (v. 3.44.3) [92]. Moreover, due to the large size of the datasets, an automatic block-wise network construction and module detection approach was chosen [93].

First, genes with zero variance across all samples were flagged and excluded. Then, for each filtered dataset a correlation matrix was calculated based on biweight midcorrelation (bicor) values and raised to specific soft thresholding powers ( ; brain: 8; heart: 10; liver: 7; muscle: 6; pancreas: 7; Supplemental Figure S5). In the cases where the scale-free topology fit index failed to reach values above 0.8, the soft-threshold power was chosen based on the number of samples [94]. The resultant signed adjacency matrices were used to compute measures of topological overlap between each pair of genes, present in Topological Overlap Matrices (TOM). Next, genes in each dataset were hierarchical clustered (average linkage method) based on topological overlap dissimilarity (1- TOM). Modules of co-expressed genes were constructed accounting for a minimum size of 50 genes, a dendrogram branch merge cut height of 0.15, and default module detection sensitivity (*deepsplit* = 2) for all datasets except for the liver (*deepsplit* = 4).

### Identification of age-associated modules, hub genes, and DEG-module-hub overlapping genes

An initial selection of modules was based on the association of each module eigengene (ME) with aging. ME is the first principal component of the expression matrix of a module and is usually considered to be the most representative gene expression profile of that group of correlated genes. The association between a given module and the trait of interest was calculated using bicor values. All modules whose ME displays a significant (FDR adjusted *p-values* < 0.05), moderate or higher (≥ 0.4) correlation with age were selected for subsequent analyses. Next, for each of the selected modules, module membership (MM) and gene significance (GS) measures were calculated. MM results from correlating the expression of individual genes to the ME, whereas GS corresponds to the absolute value of the correlation between individual genes and the trait of interest. Similar to the previous step, only modules with moderate or higher (≥ 0.4) and significant (*p-values* < 0.05) correlations were considered to be relevant. Lastly, for each selected module, genes with individual GS > 0.2 and MM > 0.8 were considered to be the most functionally important, i.e. hub genes (as seen in [95–97]). The R package UpSetR (v. 1.4.0) [98] was used to calculate and visualize the overlap between the DEGs, module genes, and hub genes.

### Functional characterization of DEG-module-hub genes’ overlap

To functionally characterize the gene lists corresponding to the intersection of DEG, module, and hub genes per tissue, we performed over-representation analysis of GO BPs using the R package clusterProfiler (v. 3.16.0) [99]. Because this package requires NCBI’s Entrez Gene IDs as input, we converted gene symbols into EntrezIDs with the org.Mm.eg.db R package (v. 3.11.4) [100]. GO terms with an FDR adjusted *p-value* less than 0.05 were selected for subsequent analyses.

### Network visualization of functionally enriched terms

Network visualization of the enriched GO terms used the enrichmentMap plugin (v. 3.3.0) [101] of Cytoscape (v. 3.8.0) [102], with nodes representing GO terms, and edges depicting similarity scores based on the number of genes in common between nodes. To construct our networks, we set an edge similarity cutoff of 0.7. GO term redundancy was addressed with the AutoAnnotate (v. 1.3.3) [43], clusterMaker2 (v. 1.3.1) [103], and WordCloud (v. 3.1.3) [104]. Similar GO terms were clustered together using the Markov Clustering Algorithm (MCL), also with an edge similarity cutoff of 0.7, and cluster labels were created with the default label algorithm Adjacent Words, with 4 maximum words per label and an adjacent word bonus of 8.

## Author Contributions

M.F. carried out the RNASeq data analysis and the gene network analysis, participated in the definition of the questions and wrote the paper; S.F. wrote and corrected the paper; M.P. and A.R. developed pipelines for data analysis; A.N. and A.R.S. corrected the paper; A.J.R., N.S. and G.M. contributed to the definition of the experimental work plan; and M.A.S.S. coordinated the overall project, defined the questions and defined the work plan, wrote and corrected the paper.

## Supporting information

Supplemental Discussion

## Acknowledgments

We are most thankful to the University of Aveiro Genome Medicine Laboratory and Institute of Biomedicine - iBiMED for supporting this work.

## Abbreviations

*Abcg2*: ATP binding cassette subfamily G member 2 (Junior blood group)
*Acaa2*: acetyl-Coenzyme A acyltransferase 2 (mitochondrial 3-oxoacyl-Coenzyme A thiolase)
*Acsl1*: acyl-CoA synthetase long-chain family member 1
*AD*: Alzheimeŕs Disease
*Anxa1*: Annexin A1
*Anxa2*: Annexin A2
*Apeglm*: Approximate Posterior Estimation for generalized linear model method
*Aβ*: Amyloid-β
*B2m*: beta 2 microglobulin
*Bicor*: Biweight Midcorrelation
*Bmp7*: bone morphogenetic protein 7
*BP*: Biological Process
*C3*: complement component 3
*C6*: complement component 6
*C8b*: complement component 8 beta polypeptide
*C9*: complement component 9
*Cdh1*: cadherin 1
*Ctsd*: cathepsin D
*Ctsh*: cathepsin H
*Dap*: death-associated protein
*DEG*: Differentially Expressed Gene
*Dlat*: dihydrolipoamide S-acetyltransferase (E2 component of pyruvate dehydrogenase complex)
*vDnaja3*: DnaJ heat shock protein family (Hsp40)
*Fasn*: fatty acid synthetase
*Fbxo8*: F-box protein 8
*FC*: Fold-change
*Fcgr2b*: Fc receptor, IgG, low affinity IIb
*FDR*: False Discovery Rate
*GEO*: Gene Expression Omnibus
*GO*: Gene Ontology
*GS*: Gene Significance
*H2-D1*: histocompatibility 2, D region locus 1
*H2-K1*: histocompatibility 2, K1, K region
*H2-T23*: H-2 class I histocompatibility antigen D-37 alpha chain
*Igf1*: insulin-like growth factor 1
*LBD*: Lewy Body Dementia
*log2FC*: log2 Fold-change
*Lrp1*: low density lipoprotein receptor-related protein 1
*ME*: Module Eigengene
*MHCI*: Major Histocompatibility Complex I
*MM*: Module Membership
*PCA*: Principal Component Analysis
*Pdha1*: pyruvate dehydrogenase E1 alpha 1
*Pdhb*: pyruvate dehydrogenase (lipoamide) beta
*Pdk2*: pyruvate dehydrogenase kinase, isoenzyme 2
*Psmb8*: proteasome (prosome, macropain) subunit, beta type 8 (large multifunctional peptidase 7)
*Pstpip2*: proline-serine-threonine phosphatase-interacting protein 2
*RNA-Seq*: RNA Sequencing
*Serpina3k*: serine (or cysteine) peptidase inhibitor, clade A, member 3K
*Serpina3n*: serine (or cysteine) peptidase inhibitor, clade A, member 3N
*Serpinf1*: serine (or cysteine) peptidase inhibitor, clade F, member 1
*Serping1*: serine (or cysteine) peptidase inhibitor, clade F, member 1
*s-IBM*: Sporadic Inclusion-body Myositis
*TDEG*: Top Differentially Expressed Gene
*TK*: tissue kallikrein
*TOM*: Topological Overlap Matrices
*VST*: Variance Stabilizing Transformation
*WGCNA*: Weighted Gene Correlation Network Analysis
*z.K*: Standardized Connectivity

