## Supplementary figures and images for "Integration of differential gene expression with weighted gene correlation network analysis identifies genes whose expression is remodeled throughout physiological aging in mouse tissues"

### Supplemental Figure 1

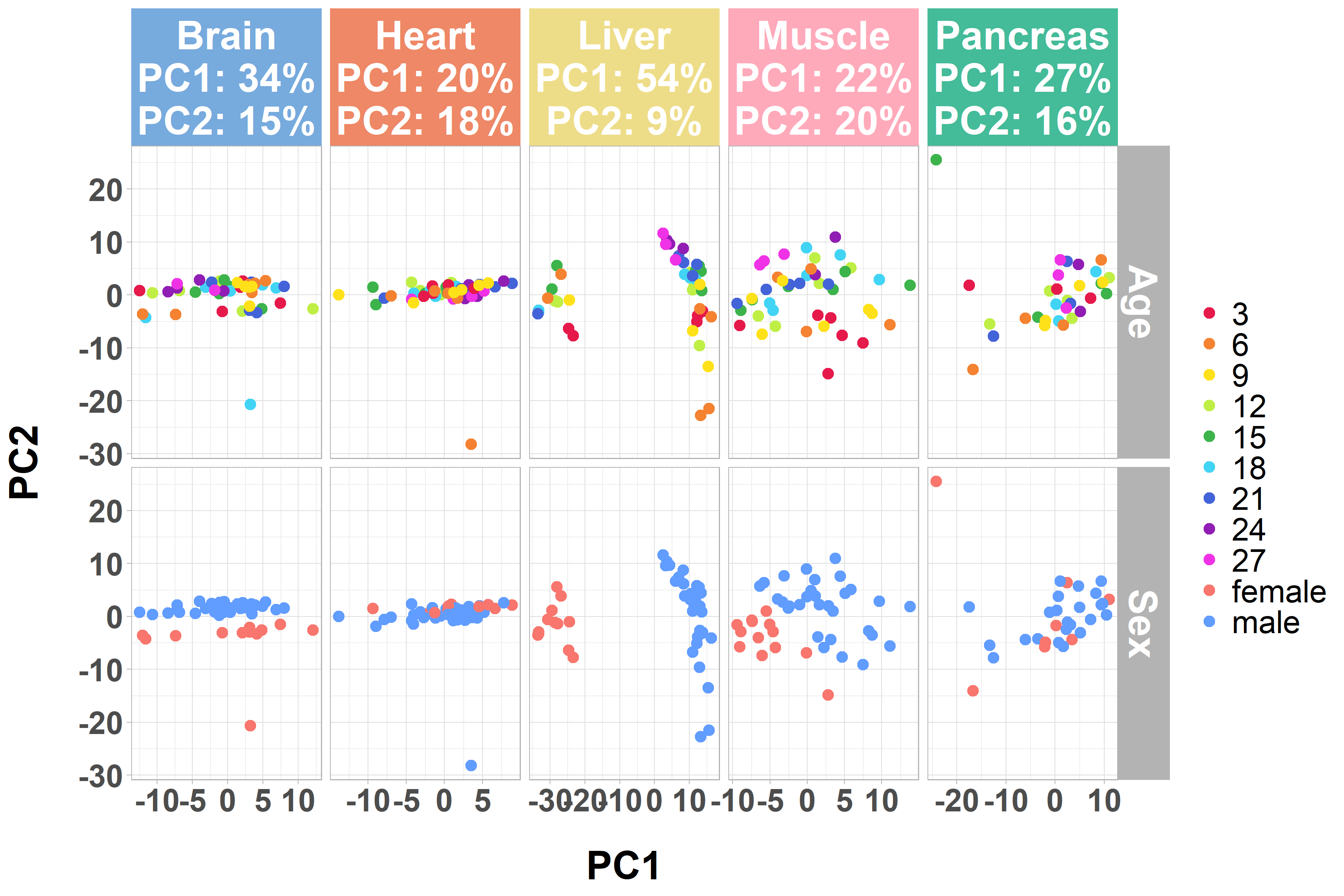

### Supplemental Figure 2

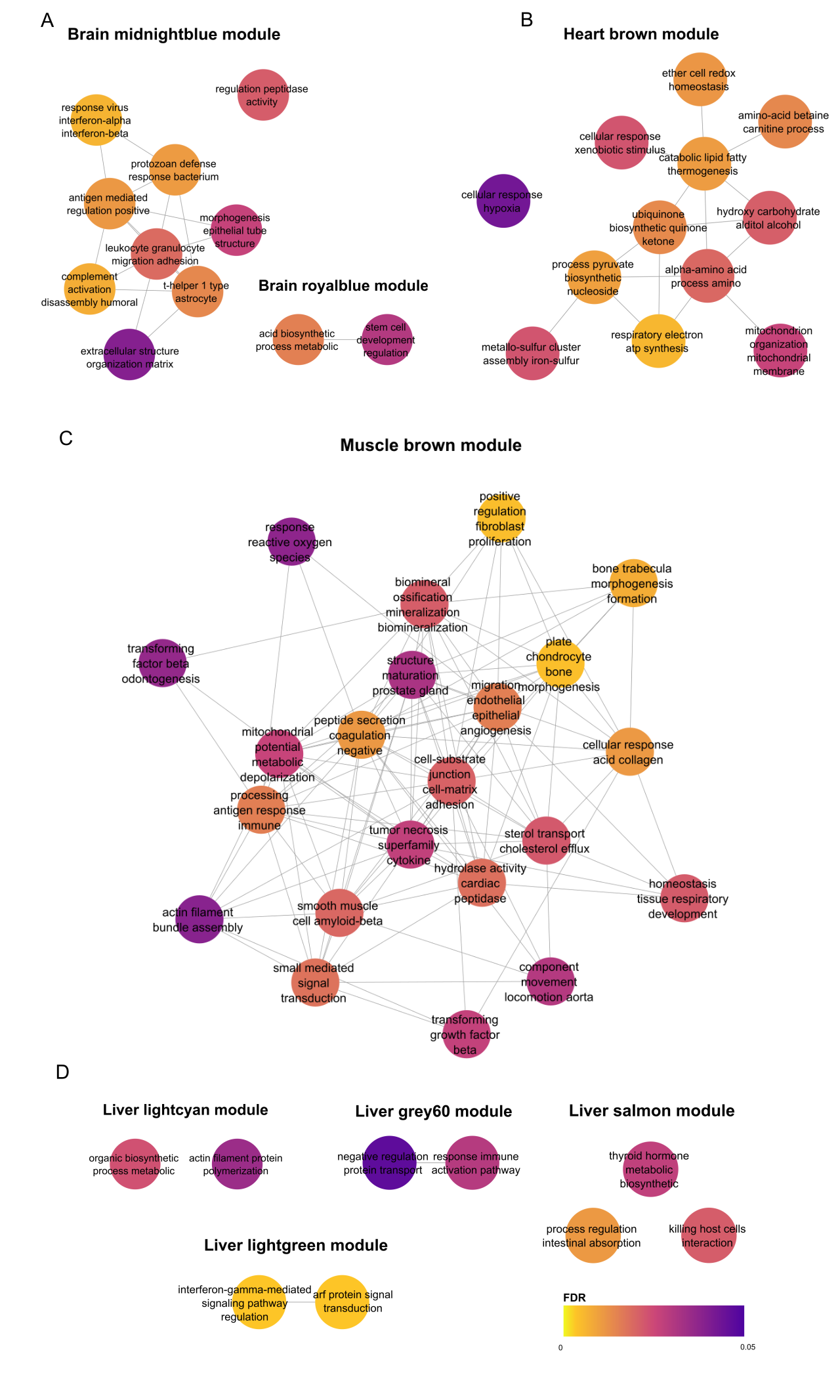

### Supplemental Figure 3

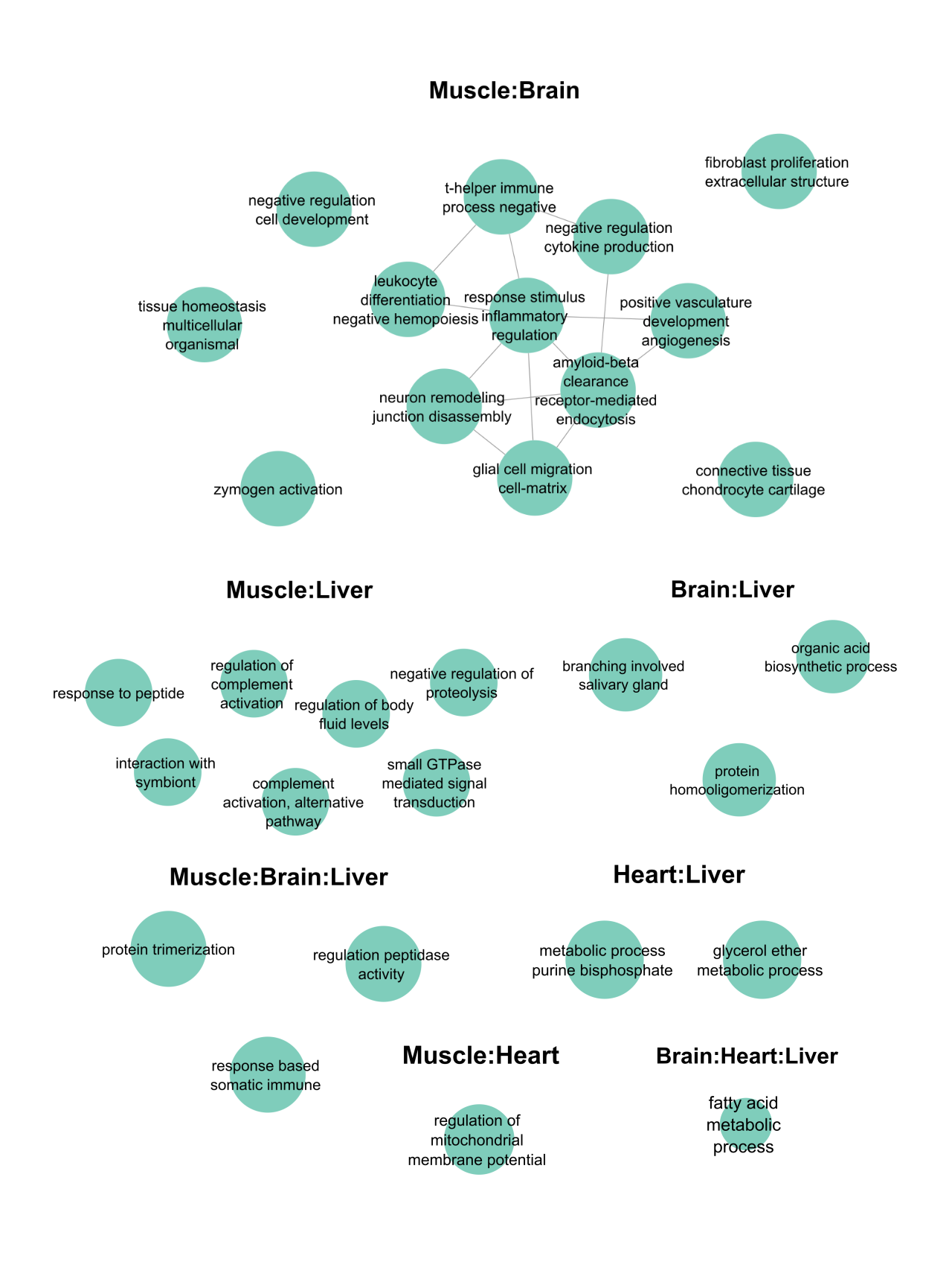

### Supplemental Figure 4

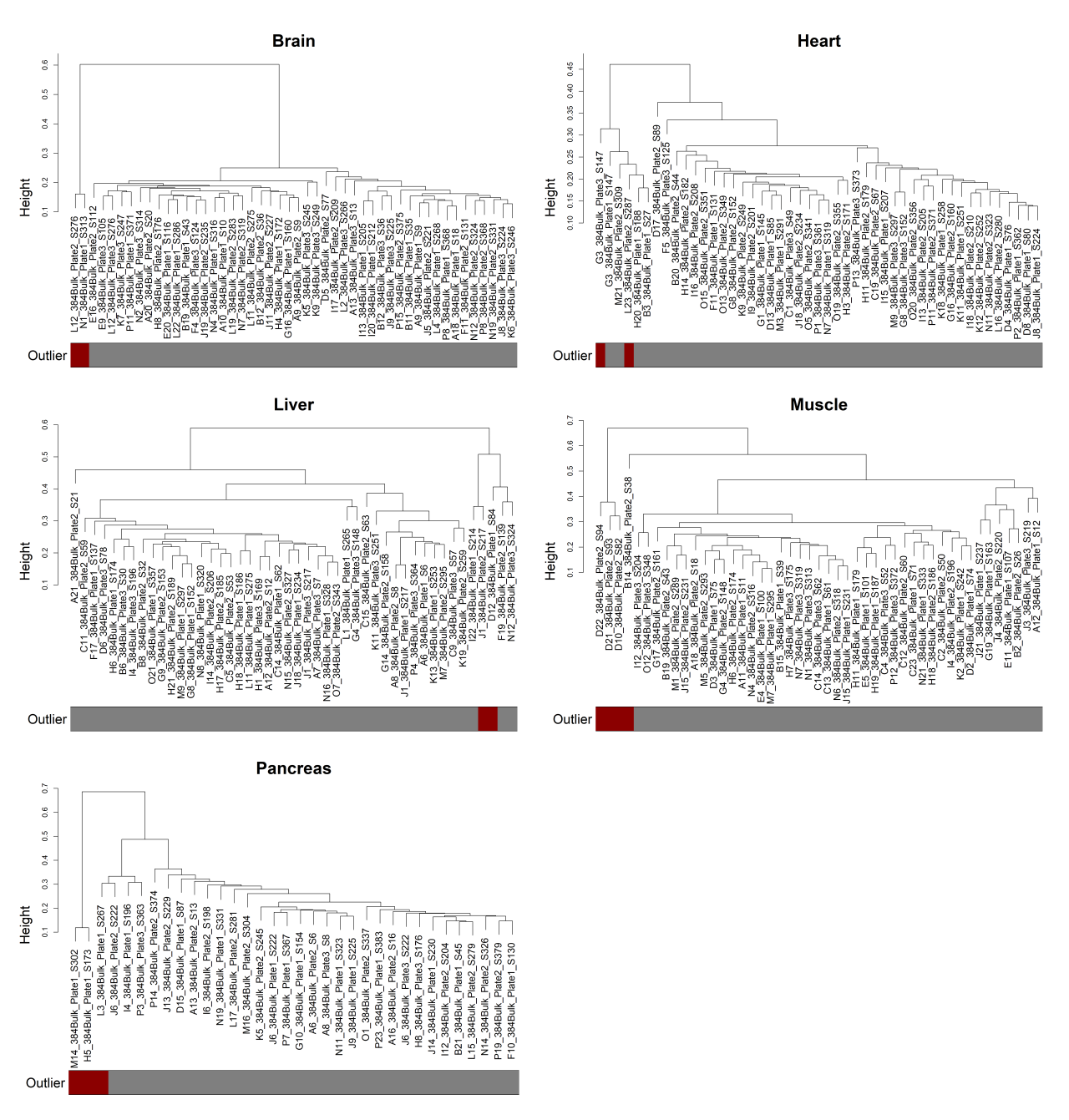

### Supplemental Figure 5

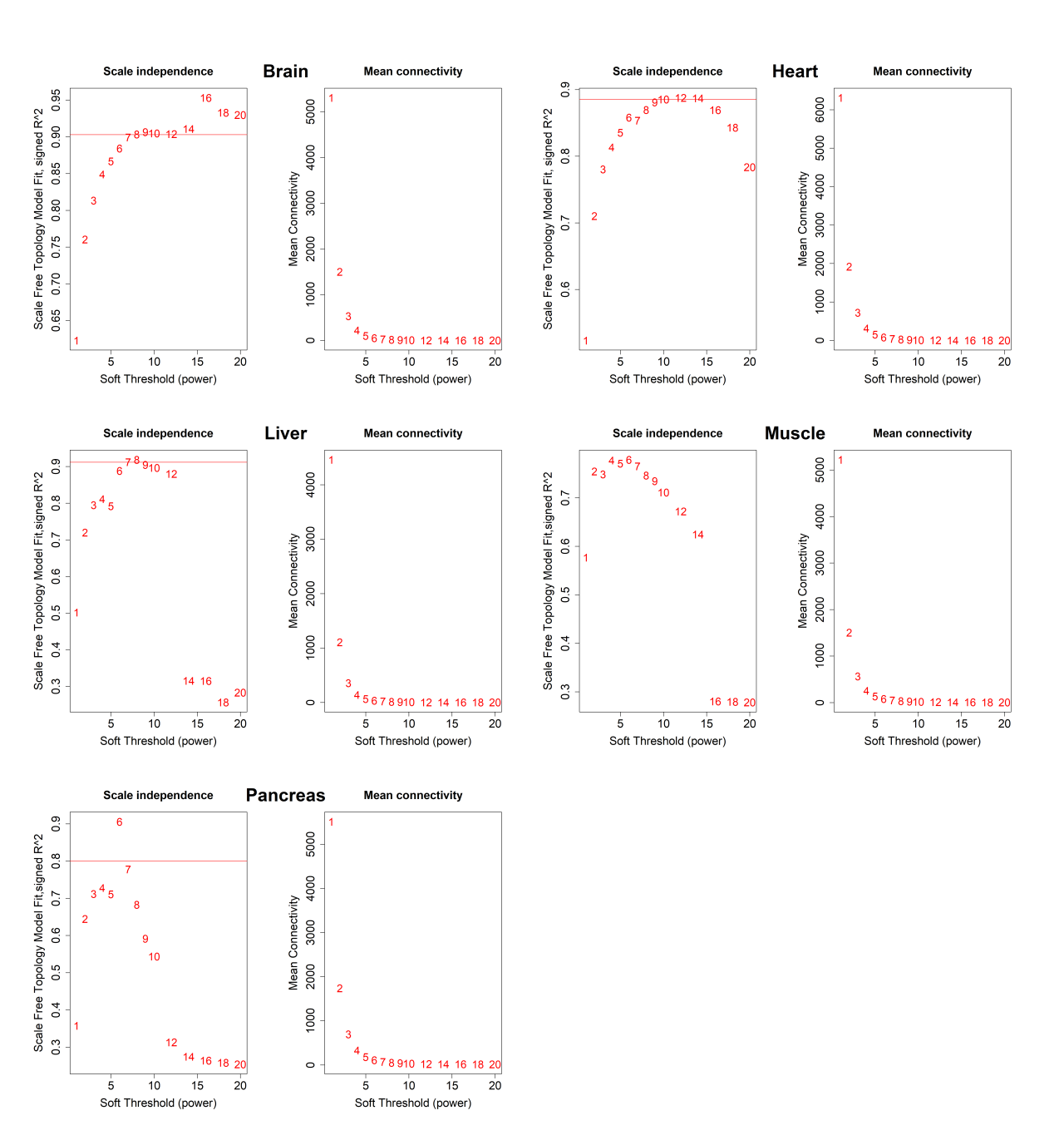
