## Supplemental Discussion for "Integration of differential gene expression with weighted gene correlation network analysis identifies genes whose expression is remodeled throughout physiological aging in mouse tissues"

### *Lipids may play an important role in organismal aging*

Notably, lipid transport and metabolism may be important in inter-tissue aging due to at least one TDEG related to these processes being identified in 4 of the 5 studied tissues. The fact that these genes are differentially expressed relatively early in the lifespan and continue to exhibit altered expression until old age may suggest a vital role of lipids in organismal aging.

In the brain, we observed a pervasive upregulation of the *Abca8a* gene, coding for an ATP-binding cassette (ABC) transporter with age. ABC transporters have been implicated in the maintenance of lipid homeostasis in this tissue, and also suggested to play a role in neurodegenerative diseases (as reviewed in 1). However, little is known about *Abca8a* and its closest human homolog *ABCA8* (*ATP Binding Cassette, Subfamily A, Member 8*) (2). The functional characterization of ABCA8 uncovered a strong and positive correlation of its mRNA expression with age and further unveiled a role in myelination and brain lipid homeostasis by regulating sphingomyelin production and cholesterol efflux in oligodendrocytes (3). Moreover, ABCA8-mediated abnormal myelination was proposed to be an early event in *Multiple System Atrophy* (MSA) (4,5). Interestingly, despite its early onset and rarity, MSA pathogenesis shares some aging hallmarks, such as proteostasis disruption with the misfolding of α-synuclein, ‘prion-like’ propagation of aberrant proteins and proteasomal impairment, as well as mitochondrial dysfunction and oxidative stress (6,7). Similarly, we found the recurrent age-related up-regulation of *Cds1* in the heart*.* This gene encodes a CDP-diacylglycerol synthase involved in the conversion of phosphatidic acid (PA) to CDP-diacylglycerol (CDP-DAG) in the endoplasmic reticulum (ER), a key intermediate to the synthesis of phosphatidylinositol (PI) in this organelle (8). PI is a membrane phospholipid that can be reversibly phosphorylated into different phosphoinositides, which are key participants in several signaling cascades thus contributing to the regulation of numerous cellular functions (9,10). In the heart, phosphoinositide signaling is most likely involved in the modulation of the electrical activity of cardiomyocytes through the regulation of ion channel activity (11). Interestingly, increased expression of *Cds1* was found in the heart of old, spontaneously hypertensive heart failure (SHHF) rats (12), as well as in vasopressin induced hypertrophic H9c2 cardiomyoblasts (13). One suggested explanation for the seemingly-indissociable increase in *Cds1* mRNA levels in the aging and diseased heart is the need to replenish PI levels after dysregulation of signaling pathways that over-consume phosphoinositides (8,12,13). Conversely, in the liver we found the *Aadac* gene to be pervasively downregulated across the lifespan. *Aadac* is a gene encoding for arylacetamide deacetylase, an ER lipase involved in hepatic triacylglycerol (TG) metabolism (14). Excessive TG accumulation in hepatocytes is one hallmark of non-alcoholic fatty liver disease (NAFLD) (15,16) and AADAC may be an important player in lipid homeostasis by regulating TG levels as its expression in rat hepatoma cells lacking endogenous AADAC was found to decrease TG accumulation (17,18). Moreover, decreased AADAC expression and activity was found in Huh7.5 cells, a hepatocyte-derived cellular carcinoma cell line, infected with the Hepatitis C virus (HCV) (19), as well as in the livers of obese patients (20), both HCV and obesity being known causes of NAFLD, a common disease among the elderly (21). Together, the existing evidence points to a role of *Aadac* gene expression in maintaining hepatic lipid homeostasis, with the observed down-regulation of its expression with aging possibly being involved in deleterious age-related TG accumulation and liver disease. Lastly, *Hacd1* also exhibited recurrent decreased expression with aging in the muscle. *Hacd1* encodes an ER enzyme involved in the synthesis of very long chain fatty acids (VLCFAs) (22,23), and has been implicated in muscle function, with mutations in this gene being reported to cause myopathy in both humans (24) and dogs (25). Moreover, HACD1 deficiency was found to impair growth and differentiation in C2C12 myoblasts (26) and to cause myofiber hypotrophy in mice and dogs (22). The exact mechanisms involved in the myopathic phenotype are not clearly understood but the link between *Hacd1* under-expression and loss of muscle function has been established and may be related to sarcopenia, the age-associated loss of muscle mass (27).

### *Tissue-specific aging expression signatures reflect an age-related loss of tissue function and homeostasis*

We also found that the most recurrently differentially expressed genes (TDEGs) in each tissue are essentially involved in tissue-specific biological processes. In the brain, protocadherins (such as *Pcdhb9*, *Pcdhga7* and *Pcdhga2*) have been implicated in neural circuit formation (recently reviewed in 28), whereas in the heart, stability and balanced expression of the sodium channel, type IV beta subunit encoded by the *Scn4b* gene have been implicated in the proper generation and conduction of the cardiac action potential (29). Another example is the *Slc22a30* gene in the liver, encoding for an organic anion transporter belonging to the SLC22 family, whose members are involved in xenobiotic transport, one of the main functions of the liver (30,31). Interestingly, in the muscle the *Sspn* gene has no biological processes annotated however existing evidence points to a role in muscle function through improved membrane stability (32,33). Similarly, in the pancreas, the *1810007D17Rik* non-coding RNA is not annotated with any biological process but was found to be largely down-regulated in cerulein-induced chronic pancreatitis in mice (34), suggesting a potential role in pancreatic homeostasis regulation.

In line with these observations, the tissue-specific aging signatures resulting from the combination of the two methodologies described in the main manuscript also reflect an age-related loss of tissue function and homeostasis. In the brain, our findings are concordant with the available literature, indicating that genes involved in immune responses, especially in MHC antigen processing and presentation, are increasingly expressed during aging in the brains of C57BL/6 mice, and probably constitute early markers of age-related neurodegeneration.  Apart from *H2-T23*, all of the MHCI genes identified have been recently shown to be upregulated with age in several regions of the central nervous system (CNS) in the mouse (35). Interestingly, higher levels of MHCI components have been linked with limited synapse density in the mouse hippocampus (36), as well as age-related synaptic loss in murine neuromuscular junctions (37), while other evidence points to an important role of MHCI in maintaining synaptic plasticity in healthy aging brain (38).  Importantly, our results are consistent with observations made in long-lived primate species and human fibroblasts that showed increased expression of MHC antigen presentation pathway genes with age, particularly *B2m* (39). Furthermore, increased expression of *B2m* has also been shown to result in impaired hippocampal neurogenesis in aged mice, thus contributing to cognitive decline (40). We also observed the upregulation of genes involved in stem cell development in the brain. *Bmp7*, a member of the TGF-β superfamily, is widely expressed throughout the adult central nervous system, and traditionally known to be involved in cortical development (41,42). Notably, *Bmp7* expression levels have been found to be elevated in the *substantia nigra* during healthy aging, only significantly decreasing following brain lesions (43), which is in line with the age-related increased expression of this gene we found. As for *Cdh1*, the protein encoded for this gene belongs to the cadherin superfamily, a group of cell-cell adhesion molecules involved in neural development (reviewed in 44,and 45). *Cdh1* has been shown to regulate neural stem cell self-renewal in adult mouse forebrain (46) and has also been implicated in proper GABAergic synapses in cultured cortical neurons (47). Moreover, altered *Cdh1* expression has been associated with tumorigenesis in the brain (48–50), as well as with blood brain barrier (BBB) permeability modulation (51,52). Taken together, our results suggest that genes involved stem cell development may be important in maintaining synaptic integrity and signaling throughout the murine lifespan. More research is needed in order to establish the role of stem cell development genes during brain aging.

Conversely, in the heart, we observed the downregulation of genes involved in energy metabolism, particularly of *Pdha1*, *Pdhb*, *Dlat*, *Pdk2*, *Acsl1* and *Acaa2*. Interestingly, *Pdha1*, *Pdhb*, *Dlat* and *Pdk2* are all involved in the irreversible oxidative decarboxylation of pyruvate, with *Pdha1*, *Pdhb* and *Dlat* encoding for the catalytic enzymes (E1 α and β subunits, and E2, respectively) of the pyruvate dehydrogenase complex (PDC), and Pdk2 encoding for a PDC regulatory enzyme (53,54). Although its role in the aging heart remains unclear, PDC is crucial in mitochondrial energy production with its end products acetyl-CoA and NADH being central molecules in the Krebs cycle and mitochondrial respiration, respectively (53,55). Higher efficiency of PDC activity has been reported in older F344 rats, mainly due to a significant decrease in the expression of regulatory pyruvate dehydrogenase kinases (PDK4 and PDK2) (56). More recently, heart failure patients reportedly showed increased PDC activity in the left ventricular myocardium, characterized both by greater expression levels of PDC catalytic enzymes, including E1α, E1β, and E2, and also by the decreased expression of PDK4, but not of PDK2 (57). It has also been shown that PDC activation is able to improve cardiac function in murine hearts (58), with the beneficial effects of PDC activity on heart function probably being due to increasing energy production under large energetic demand conditions. Furthermore, Acsl1 and Acaa2 are both involved in fatty acid beta-oxidation, which also generates acetyl-CoA and NADH (59–61). Despite not being studied in the context of aging, both genes have been shown to play a role in maintaining proper cardiac function, with possible implications for age-associated heart dysfunction. In fact, it was recently shown that over-expression of ACSL1 reduced cardiac hypertrophy and dysfunction (62). More indirectly, in heart failure-induced rats, a treatment successfully improved myocardial energy metabolism through the upregulation of the expression of genes involved in fatty acid metabolism, including Acaa2 (63). Additionally, an aging-induced decline in fatty acid oxidation has also been reported in the hearts of aging mice (64). Integrating these findings with our observations of decreased expression of genes involved in pyruvate and fatty acid β-oxidation, indicates a general impairment of cardiac energy substrate metabolism, and suggests that the energy requirements of the aging heart are severely compromised.

We also found a general decline in the expression of the genes comprising the muscle-specific aging signatures. Unsurprisingly, these genes are mainly involved in regulating muscle hypertrophy, regeneration, and homeostasis, as is the case of *Igf1* and *Lrp1*. *Igf1* is a paramount growth factor that is released in this tissue during exercise, activating the Akt/Protein Kinase B–mTOR pathway, thus promoting ribosomal biogenesis and translation to produce new myofibril proteins (65). Under normal conditions of rest, circulating Igf-1 protein levels are significantly greater in young adults compared to older individuals (66). In fact, throughout aging, the skeletal muscle’s ability to induce protein synthesis in response to protein ingestion and exercise declines, leading to sarcopenia, the age-related phenotype of reduced muscle mass and function, suggesting a reduced IGF-1/phosphatidylinositol 3-kinase (PI3K)/Akt signaling with aging in this tissue (67,68). Moreover, overexpression of local skeletal muscle Igf1 isoforms, namely Igf-1Ea and Igf-1Eb, has been shown to stimulate muscle hypertrophy (69). Regarding *Annexin A1* and *A2*, these genes encode for proteins belonging to the annexin family and are known to play an important role in plasma membrane repair of skeletal muscle cells (70–72). Loss of *Anxa1* has been shown to affect the fusion of muscle satellite stem cells, thus delaying muscle regeneration (73), while loss of *Anxa2* is associated with impaired myofiber repair and regeneration as well as progressive muscle weakening with age (74). Interestingly, and contrary to our observations, the expression of both genes has been found to increase with age both in healthy humans (75) and in *ad libitum* fed rats (76). Moreover, *Lrp1* is a large endocytic receptor involved in muscle fibrosis, where Lrp1-Decorin pathway leads to activation of TGF-β, promoting the expression of pro-fibrotic molecules (77,78). Additionally, *Lrp1* depletion impairs fracture repair in the bones of old mice, while overexpression improves it (79). Notwithstanding these observations, the role of *Lrp1* should be further elucidated in the context of skeletal muscle aging. Overall, our results indicate that skeletal muscle structure and functioning declines with age, where key genetic players in muscle regeneration are found to be downregulated. However, the exact role and underlying mechanisms of these genes in the aging mammalian skeletal muscle remains a marker of interest to be explored in future studies.

Lastly, we observed an overall decrease in expression of genes during hepatic aging, with significantly dysregulated genes including *C6*, *C8b*, *C9*, *Cdh1*, *Dnaja3* and *Abcg2*. These genes are involved in a plethora of functions pivotal in maintaining homeostasis both at the tissue- and organismal-level. The liver produces around 90% of complement proteins as the complement system plays a pivotal role in liver homeostasis and immune responses (80,81) while diminished levels of soluble complement components are found in cirrhosis and liver failure (81). In line with our observations, *C6*, *C8b* and *C9* have been reported as being downregulated in the livers of old mice (82). Interestingly, the identified genes are members of the membrane attack complex (MAC) (83), suggesting an age-related impairment of complement activation at the level of MAC assembly. *Abcg2* is an ABC transporter mainly expressed in the liver, intestine and kidney, and known to play an important role in mediating xenobiotic absorption and excretion (84,85). Notwithstanding these protective mechanisms, and indeed because of them, *Abcg2* has also been associated with multidrug resistance (reviewed in 86, also addressed in 87). Moreover, altered expression of this transporter has also been reported in hepatic disease such as Human Obstructive Cholestasis (88) and Hepatocellular Carcinoma (HCC) (89,90). Importantly, *Abcg2* expression has been reported to decrease in the aging murine liver (91–93), and in humans (94). These findings are in line with our observations of a progressively decreased expression of this gene across the mouse lifespan and corroborate the hypothesis of age-related compromised detoxification processes (93,95). Furthermore, the *Cdh1* gene encodes for E-cadherin, a cell-to-cell adhesion molecule expressed by hepatocytes and biliary epithelial cells in the liver (96). *Cdh1* is critical in maintaining liver homeostasis and suppressing carcinogenesis, and loss of *Cdh1* in intrahepatic biliary epithelial cells has been shown to promote fibrosis as well as cancer progression due to impairment of the intrahepatic biliary network function and inflammatory responses that lead to progenitor cell proliferation (96). Despite being annotated as involved in the positive regulation of protein import into the nucleus (97), this role was inferred from sequence orthology and still needs further experimental confirmation. Consistent with our result, *Cdh1* was also found to have a decrease in translation output and efficiency with aging (30). In addition, *Dnaja3* gene encodes for a member of the DNAJ/Heat shock protein 40 (Hsp40) family, known to stimulate heat shock protein 70 (Hsp70) activity through ATP hydrolysis, enabling substrate delivery to Hsp70 (98). Interestingly, Hsp70 has been proposed to be involved in the early phase of liver regeneration by the induction of tumor necrosis factor alpha (TNF‐α) (99), and our observations of decreased expression of *Dnaja3* with age may suggest an age-related impairment in hepatic regeneration through diminished Hsp70 activity. However, further studies are required to elucidate the role of *Dnaja3* in the aging liver. Conversely, *Pstpip2* was found to be upregulated in our results. This gene may be an important marker and/or target for hepatic fibrosis as, in liver macrophages, it has been found to be hypermethylated while its overexpression reduced inflammation and alleviated hepatic fibrosis in mice (133). Taken together, these results evince an overall functional dysregulation in the aging mouse liver with particular emphasis placed once again on the genes involved in immune response and cell-cell adhesion.

### *Dysregulation of organic acid metabolism across the lifespan is shared by the brain, the heart and the liver*

As for the brain, heart and liver, the biological processes commonly affected by aging relate to organic acid metabolism.

In the brain, we report an age-related increase in expression of some genes implicated in the synthesis of prostaglandins (PGs) from the fatty acid precursor arachidonic acid, namely *Aldh1a2*, *Ptgds*, and *Sphk1*. Interestingly, in this tissue PGs have been linked to neuroinflammation and associated memory loss during aging and in age-related diseases (100–102). *Sphk1*, encodes for an enzyme responsible for catalyzing the phosphorylation of sphingosine into sphingosine-1-phosphate (S1P), a role that makes this gene very relevant in the context of aging and neurodegeneration, due to growing evidence demonstrating the activation of the IGF-I-Akt-mTOR pathway by S1P (recently reviewed in 103). Decreased levels of S1P and Sphk1 activity have been reported in AD pathogenesis (104), suggesting that our observations of increased *Sphk1* expression in the aging brain might be a compensatory mechanism of loss of Sphk1 function. Moreover, *Ptgds* encodes a protein responsible for the synthesis of prostaglandin D2 (PGD2), playing an important neuroprotective role in AD as an Aβ chaperone (105–107), while also being linked with several other brain disorders (reviewed in 101). Furthermore, *Aldh1a2*, involved in the retinoid signaling pathway, is responsible for catalyzing the conversion of retinaldehyde to retinoic acid in the brain (reviewed in 108). Interestingly, early decline of retinoic acid signaling has been implicated in animal models of AD and frontotemporal dementia (109). *Aldh1a2* has also been implicated in regulating the formation, proliferation and differentiation of neural stem cells while also regenerating ventral motor neurons (110). Additionally, a significant decrease and redistribution of ALDH1A2 in adult spinal cord has been linked with neuronal dysregulation and death in Amyotrophic Lateral Sclerosis (111). Curiously, the synthesis of prostaglandins and retinoic acid in the brain has also been suggested to play a role in senescence, as the levels of both *Ptgds* and *Aldh1a2* mRNAs were found to be significantly up-regulated in the hippocampus of senescence-accelerated prone (SAMP8) mice, when compared to senescence-accelerated resistant (SAMR1) controls (112). Overall, our observations of increasing levels of genes involved in PG synthesis in older mice are consistent with the existing evidence.

In the liver, we observed an age-associated increase in the expression of *Fasn* which encodes fatty acid synthetase, an enzyme known to catalyze the synthesis of long-chain fatty acids, hence playing an essential role in *de-novo* lipogenesis (113). In line with our results, a recent study has linked fatty acid synthase activity with the induction of senescence where increased *Fasn* expression was reported in senescent-induced hepatic stellate cells (HSCs), as well as in liver samples from aged mice when compared to younger individuals (113), suggesting heightened fatty acid synthesis with age. Moreover, FASN inhibition was sufficient to halt the induction of senescence in HSCs, as observed by the prevention of the senescent-induced cell cycle arrest, as well as of the up-regulation of different markers of senescence (113). Age-related increase in *Fasn* expression has also been linked to a lipogenic switch from glycolysis to mitochondrial respiration (114,115). Furthermore, in NAFLD mouse models and patients, there are significant correlations between *Fasn* mRNA and protein expression with hepatocellular lipid accumulation and steatosis in the liver (116). Furthermore, this enzyme may also be implicated in purine metabolism. Purines are nitrogen-containing heterocyclic molecules of key importance in cellular homeostasis as they are involved in a plethora of processes, such as the synthesis of nucleic acids, and the regulation of energy, lipid and carbohydrate metabolism, among others (117,118). In the liver, the association between aging and purine metabolism dysregulation is tentative, with most evidence supported by the central role of uric acid (UA) in age-related liver diseases. UA is the final product of the *de novo* purine biosynthetic pathway (117,118), and is now a known regulator of lipid metabolism (reviewed in 119, also addressed in 120), with existing evidence pointing to an important role of UA in the development of metabolic diseases. In fact, high levels of UA in human hepatocellular carcinoma (HCC) cells (HepG2 cells) have been associated with an oxidative stress-mediated increase in *de novo* lipogenesis and triglyceride accumulation, important indicators of hepatic steatosis development (121). Moreover, in the same cell line, UA was found to directly induce the expression of lipogenic enzymes such as acetyl-CoA carboxylase 1 (ACC1), stearoyl-CoA desaturase 1 (SCD1), and fatty acid synthase (FAS) (122). Interestingly, despite not evaluating the role of UA in the tumorous phenotype, dysregulated purine metabolism was also observed in HCC patient samples, particularly the up-regulation of key enzymes responsible for catalyzing purine biosynthesis, and the reduction of tumor proliferation resulting from the targeted ablation of the purine *de novo* synthesis pathway (123). Overall, these findings are in line with our observation of increased expression of *Fasn* with age in the liver, although more studies are required to clarify the role of impaired purine metabolism in hepatic aging and to draw more definite conclusions.

Remarkably, alterations in purine metabolism have also been observed in the hearts of old and senescent mice, particularly an increased extracellular accumulation of inosine monophosphate (IMP), hypoxanthine, xanthine and uric acid, central players in the purine biosynthetic pathway (117,124). Conversely, we observed a down-regulation of adenosine kinase (*Adk*) in old mice, starting in the transition from 18 to 21 months (Table 3; Supplementary Table S6). This enzyme is involved in the purine salvage pathway through the phosphorylation of adenosine back to adenosine monophosphate (AMP) (117). Purine metabolism has been implicated in disease-associated cardiac dysfunction, as in mouse models of Huntington’s disease (HD) some genes involved in *de novo* purine biosynthesis were found to be significantly down-regulated (125). Interestingly, the authors also observed decreased levels of adenosine triphosphate (ATP), accompanied by a 2-fold down-regulation of adenylate kinase 1 (*Ak1*), an enzyme important in the inter-conversion of adenine nucleotides (125,126). In line with these findings, we found *Ak1* to be downregulated in the hearts of old mice (Table 3; Supplementary Table S6). Together, these results highlight the role of impaired purine metabolism in heart dysfunction, with adenine metabolism potentially playing an important part.

In addition, a decrease in expression of genes involved in fatty acid oxidation was also observed in the heart, particularly of *Acaa1a*, *Acaa2*, *Acadl*, *Adipor1*, *Auh*, *Cpt*, *Crat*, *Eci2*, *Etfb*, and *Hadh*, which corroborates previous reports of age-related cardiac dysfunction mediated by cardiac lipotoxicity as a result of impaired oxidation of fatty acids (recently reviewed in 127).

### *Age affects mitochondrial membrane potential in both the heart and the muscle*

The muscle and the heart share an age-related dysregulation of mitochondrial membrane potential, which is not surprising as mitochondrial dysfunction has long been associated with aging, being that loss of membrane potential (ΔΨm) is one of the hallmarks of aging in this organelle (128,129). In the muscle, the genes involved in this process are *Dcn* (*Decorin*), *Myoc* (*Myocilin*), and *Pid1* (*Phosphotyrosine Interaction Domain Containing 1*), and despite the lack of evidence regarding their role in this tissue, studies in various cell types and organs associate their expression with the loss of mitochondrial membrane potential. Decorin-induced inhibition of ΔΨm has been reported in breast carcinoma cells (130) and human umbilical vein endothelial cells (HUVEC cells) (131). In human trabecular meshwork cells, myocilin imported to mitochondria was targeted to membranes and impaired ΔΨm (132). Moreover, over-expression of *Pid1* has been found to depolarize  ΔΨm in human medulloblastoma cells (133,134), and in embryonic murine adipocytes (135–137). Interestingly, in the muscle we observed a decreased expression of these genes with aging starting around middle age, which may be suggestive of a protective mechanism of mitochondrial function with increasing age in this tissue. Nevertheless, the role of these genes in ΔΨm needs to be further addressed and clarified. In the heart, the evidence is also scarce and establishing a link between *Got1* (*glutamic-oxaloacetic transaminase 1, soluble*), *Ndufs1* (*NADH:ubiquinone oxidoreductase core subunit S1*) and *Prdx3* (*peroxiredoxin 3*), and age-related dysregulation of ΔΨm in this tissue is not straightforward. Increased expression of *Got1* induced by prostaglandin E2 (PGD2) resulted in diminished ΔΨm in mouse bone marrow macrophages (138), while Similarly, decreased ΔΨm was also exhibited by WEHI7.2 mouse thymoma cells over-expressing *Prdx3* (139). Conversely, there is more direct evidence regarding the role of *Ndufs1* on mitochondrial function in the heart. *Ndufs1* is the largest subunit of the respiratory complex I (140) and its impaired translocation to mitochondria has been shown to exacerbate the diabetic cardiomyopathy phenotype in mice through mitochondrial dysfunction, including decreased ΔΨm levels (141). Our observations of decreased expression of these genes with aging are only partially concordant with these findings. On one hand, decreased expression of *Ndufs1* may compromise the function of complex I in mitochondrial respiration, resulting in loss of ΔΨm, which is in line with the observations in mice with diabetic cardiomyopathy. On the other hand, the decreased expression of *Got1* and *Prdx3* may serve as a compensatory response against diminished ΔΨm. Similar to the muscle, more studies are needed to better understand the mechanisms behind mitochondrial membrane potential dysregulation and its potential implications for the aging heart.

Supplemental Discussion References

16.	Malhi H, Gores GJ. Molecular Mechanisms of Lipotoxicity in Nonalcoholic Fatty Liver Disease. 2008;

57.	Sheeran FL, Angerosa J, Liaw NY, Cheung MM, Pepe S. Adaptations in protein expression and regulated activity of pyruvate dehydrogenase multienzyme complex in human systolic heart failure. Oxid Med Cell Longev. 2019;2019.
