## Supplemental Legends for "Integration of differential gene expression with weighted gene correlation network analysis identifies genes whose expression is remodeled throughout physiological aging in mouse tissues"

Supplemental Figures

**Supplemental Figure S1. Transcript variance per tissue of interest after outlier removal.** PCA of selected tissues based on the top 500 most variable genes and colored by age and sex. Because sex is a considerable contributor to sample segregation it was included in the design matrix as a co-variable.

**Supplemental Figure S2. Summary network visualization of GO enrichment analysis based on the DEG-module-hub gene overlap in the brain (A), heart (B), muscle (C), and liver (D) modules.** Each *meta-node* represents a cluster of similar GO terms (colored by FDR adjusted *p-value*) and each *meta-edge* depicts genes shared between the nodes. GO enrichment analysis of selected gene sets was performed using clusterProfiler (FDR < 0.05) and enrichment maps of the obtained lists of GO terms were constructed using the EnrichmentMap plugin in Cytoscape (FDR < 0.05 and edge similarity > 0.7). Redundancy was overcome by clustering together and annotating similar terms based on the most frequent words (AutoAnnotate, clusterMaker2, and WordCloud plugins in Cytoscape; clustering algorithm: Markov cluster algorithm - MCL; labeling algorithm: adjacent words with a maximum 4 words per label and an adjacent word bonus of 8).

**Supplemental Figure S3. Summary network visualization of the GO term overlap between tissues.** Each meta-node represents a cluster of overlapping GO terms and each meta-edge depicts shared genes.

**Supplemental Figure S4. Outlier detection by sample network approach** [1]**.** Sample dendrograms were produced by hierarchical clustering using average linkage as the clustering method and 1-A (network adjacency) as the distance between samples. Samples were considered outliers (depicted in red) if their standardized connectivities (z.K, see [1]) were more than 2 standard deviations away from the mean z.K.

**Supplemental Figure S5. Soft-thresholding power determination.** Analysis of scale-free fit indexes (left panels) and mean connectivities (right panels) for different soft-thresholding powers (x-axes; red numbers). In cases where there is a lack of fit of scale-free topology, soft-thresholding powers were chosen based on sample number, as proposed by the authors [2].

Supplemental Files

**Supplemental File S1. Differentially expressed genes per tissue across the evaluated time points.** Genes with *S*-values less than 0.005 were selected. Relates to Figure 1 and to Table 1.

**Supplemental File S2. Global characterization of WGCNA modules and selected modules’ genes, with respective gene significance and module membership values.** Hub genes were considered based on module membership (MM) and gene significance (GS) higher than 0.8 and 0.2, respectively. Relates to Figure 2.

**Supplemental File S3. Gene overlap between DEGs, TDEGs, module genes and hub genes.** Genes considered were in common, at least, in the DEG, module, and hub gene sets. Relates to Figure 3 and to the left panel of Figure 4.

**Supplemental File S4. Gene Ontology (GO) over-representation and network analysis.** GO terms with FDR adjusted *p-values* less than 0.05 were considered for analysis. Relates to Figure S2 and to Table 2.

**Supplemental File S5. Gene Ontology (GO) term overlap between the brain, heart, liver, and muscle.** GO terms with FDR adjusted *p-values* less than 0.05 were considered for analysis. Relates to the right panel of Figure 4, to Figure S3 and to Table 3.

**Supplemental File S6. Sample characterization.** Number of male and female samples from the selected time points before and after the removal of low coverage and outlier samples.

Supplemental Material References

1.	Oldham MC, Langfelder P, Horvath S. Network methods for describing sample relationships in genomic datasets: application to Huntington’s disease. BMC Syst Biol [Internet]. BioMed Central; 2012 [cited 2020 Jul 16]; 6: 63. Available from: http://bmcsystbiol.biomedcentral.com/articles/10.1186/1752-0509-6-63

2.	WGCNA package: Frequently Asked Questions [Internet]. [cited 2020 Aug 16]. Available from: https://horvath.genetics.ucla.edu/html/CoexpressionNetwork/Rpackages/WGCNA/faq.html
